# From margins to mainstream: restoration of riparian margins to support oil palm sustainability in modern and future climates

**DOI:** 10.64898/2026.10.09.757517

**Authors:** Sacchi Shin-Clayton, Anak Agung Ketut Aryawan, Rebecca A. Senior, Andrea Manica, Damayanti Buchori, Jean-Pierre Caliman, Purnama Hidayat, Michela Leonardi, Mohammad Naim, Sarah H. Luke, Edgar C. Turner

## Abstract

Palm oil is the world’s leading vegetable oil, with production continuing to expand, particularly across Southeast Asia. Within established plantations, conserving and restoring riparian margins can reduce erosion, improve connectivity, and support biodiversity. Simultaneously, climate change is increasing thermal stress within tropical agricultural landscapes, raising the potential importance of riparian buffers as microclimatic refugia, although this has not previously been evaluated. The Riparian Ecosystem Restoration in Tropical Agriculture (RERTA) Project is a long-term Before–After Control–Impact experiment established within oil palm plantations in Riau, Indonesia, to evaluate alternative riparian restoration strategies following plantation replanting. Treatments included passive restoration (retaining mature oil palms and allowing natural regeneration), active restoration (enrichment planting with six native tree species), and a control (replanting immature oil palms to the river edge). Six years after restoration, we assessed vegetation structure using terrestrial laser scanning and thermal stability using temperature loggers and thermal imaging. We also evaluated the long-term temperature buffering capacity of each treatment under current and projected climates using CHELSA climate data and downscaled outputs from multiple General Circulation Models under Shared Socioeconomic Pathway scenarios for 2045 and 2085. Active restoration produced greater structural complexity, a more closed canopy, and cooler, more thermally stable conditions than passive restoration and control treatments. Restored margins also contained significantly more cool surface microhabitats, reducing thermal extremes under both current and future climate scenarios. Our findings demonstrate that active restoration of riparian margins can enhance thermal buffering and create microclimatic refugia for biodiversity within tropical oil palm landscapes. As agricultural intensification and climate change continue to reshape tropical ecosystems, restoring structurally complex riparian vegetation will be critical for sustaining biodiversity and the ecosystem services upon which agricultural production depends.

## 1. Introduction

Agriculture is one of the world’s fastest-growing industries and remains a foundation of many national economies (Dong et al., 2022; Nor Diana et al., 2022; Rehman et al., 2024). Yet, its rapid expansion and intensification are major drivers of global biodiversity loss, including declines in insect abundance and diversity, which provide essential ecosystem services and support crop production (Raven & Wagner, 2021). At the same time, agricultural production is increasingly threatened by climate change, with global temperatures projected to exceed 1.5 °C above pre-industrial levels within the 2030s, “the point of no return”, unless substantial mitigation measures are taken (FFPI, 2023; Raven & Wagner, 2021). Climate change is expected to amplify extreme weather events, including droughts, storms, and floods, further threatening ecosystem functioning and crop yields (IPCC, 2021). Many crops are highly climate-sensitive (Mihailescu and Bruno Soares, 2020), and future water stress is projected to reduce yields, lower water-use efficiency, and increase their vulnerability to pests and diseases (IPCC, 2021).

These pressures do not act in isolation. Strong synergistic effects are anticipated between agricultural intensification and climate change (Raven & Wagner, 2021; Warren et al., 2022). Insect declines, for example, have been linked to both drivers, and key functional groups such as pollinators are sensitive to changes in management and to rising temperatures due to their narrow thermal tolerance ranges (Raven and Wagner, 2021). Indeed, many tropical pollinators exhibit stress or reduced performance when temperatures exceed their optimal range by only 2–4 °C (Settele et al., 2016).

Tropical Southeast Asia is an important region to study the interactions between habitat and climate change in agricultural systems. The region represents a rapidly expanding centre of global agricultural production, particularly oil palm – the leading global source of the vegetable oil, palm oil – and is projected to experience some of the most rapid climatic changes (Outhwaite et al., 2022). Models predict a 2–4 °C increase in mean annual temperature by 2050, accompanied by altered rainfall regimes, greater rainfall seasonality, and a rising frequency of extreme weather events, such as El Niño–associated droughts and severe flooding (Appelt et al., 2023; Rehman et al., 2024). These shifts are expected to directly threaten the productivity and stability of agricultural systems in the region, while also compounding ongoing pressures from regional biodiversity loss (Zhang et al., 2024).

Within Southeast Asia, oil palm underpins the economies of several countries, with Malaysia and Indonesia alone accounting for ∼80% of global palm oil production (FAO, 2023; Vijay et al., 2016). Although oil palm achieves far higher yields per hectare than other vegetable oil crops and has relatively high ecosystem complexity and stability owing to its long replanting cycles (Meijaard et al., 2020), its expansion has been closely associated with deforestation, biodiversity loss, and ecosystem degradation (Vijay et al., 2016). Forest conversion to oil palm drives marked changes in community composition, with declines of up to 60% in species richness for key invertebrate groups. Forest specialist species across several taxa, including ants (Boyle et al., 2021; Fayle et al., 2010), bees (Lee et al., 2001), beetles (Chung et al., 2000), and termites (Luke et al., 2014), show particularly steep declines, often linked to hotter, drier microclimates and the loss of habitat complexity. In contrast, generalist and invasive species often increase in abundance in plantations (Savilaakso et al., 2014). Like other crops, oil palm is sensitive to climatic variability: it grows optimally under mean annual temperatures of 24–28 °C and annual rainfall of 1,800–2,500 mm with minimal dry-season stress, whereas prolonged exposure to temperatures above ∼33 °C, drought conditions (<100 mm per month for several months), or waterlogging can substantially reduce photosynthesis, fruit set, and yield (Corley & Tinker, 2016; Krishnan & Pimid, 2024). Furthermore, oil palm production depends on biodiversity within plantations to sustain several essential ecosystem services, including pollination, pest regulation, and soil fertility (Garbach et al., 2017; Suarez & Gwozdz, 2023), meaning that biodiversity declines associated with climate change may compound the direct effects of climatic change on yield.

One promising intervention to enhance both biodiversity and ecosystem resilience in tropical agricultural landscapes is the maintenance or restoration of riparian forest buffers. These forested zones provide essential habitat corridors and refugia for a wide range of aquatic and terrestrial species, including amphibians, birds, insects, and small mammals, thereby supporting biodiversity within intensively managed landscapes (Luke et al., 2019; Malcher et al., 2023; Wilkinson et al., 2018). Healthy and established riparian buffers are linked with positive influences on both terrestrial and aquatic ecosystems. With increased structural complexity in the stream leading to bank stabilisation, litter inputs, and climate control, which all in turn lead to improved ecological processes such as pollination, seed dispersal, and nutrient cycling, which enhance adjacent agricultural productivity through spillover effects (Garibaldi et al., 2011; Gray et al., 2015; Gray & Lewis, 2014). This, in turn, can regulate hydrology, improve water quality, and reduce erosion and agrochemical runoff into freshwater systems (Dinca et al., 2025; Riis et al., 2020). Importantly, the canopy structure of riparian areas can also reduce incoming solar radiation and lower ambient temperatures below the canopy, creating cooler and more stable microclimates (Barclay et al., 2017). This thermal buffering is especially critical for small-bodied, temperature-sensitive organisms, such as insects and amphibians, which depend on shaded refugia to mitigate heat and water stress (Ran et al., 2020).

Despite the recognised temperature regulation benefits of riparian buffers, strategies for restoring riparian forests remain poorly investigated, and their effects on temperatures are not well understood, particularly in tropical regions and under future climate scenarios. Given this variability, predicted and estimated impacts on biodiversity and biodiversity-mediated services are driving growing pressure for action and guidelines. In response to mounting global pressure to slow deforestation (EUDR 2026 Act; (European Commission, 2026), several certification initiatives and standards are already well established and have developed guidelines highlighting the importance of riparian margins and their protection from land clearing (ISPO, 2026; Roundtable on Sustainable Palm Oil, 2026).

This study developed from a long-term established partnership project with oil palm managers (the Riparian Ecosystem Restoration in Tropical Agriculture (RERTA) Project), based in established oil palm plantations in Riau, Indonesia, to evaluate the effects of passive and active restoration approaches in riparian margins on vegetation structure and thermal characteristics in the first six years of the project and under projected climate scenarios.

We address the following key questions:

1. How does vegetation structure differ across alternative restoration treatments (active replanting, passive restoration), and a control?
2. How do the temperature characteristics of an area (daily max, mean, min temperature, Vapour Pressure Deficit (VPD), and land surface temperature (LST)) differ across alternative restoration treatments?
3. How does thermal buffering capacity (temperature offset) vary across restoration treatments, both during the initial restoration years and under projected climate scenarios, with reference to published data on insect critical thermal maxima (CTmax)?

## 2. Methods

### 2.1. Study sites

The Riparian Ecosystem Restoration in Tropical Agriculture (RERTA) Project is part of the wider Biodiversity and Ecosystem Function in Tropical Agriculture (BEFTA) Programme (Luke et al., 2020), a long-term research collaboration between academia and industry. RERTA is a long-term Before-After Control-Impact (BACI) experiment, designed to evaluate the effects of alternative restoration approaches (including both passive and active) in riparian margins within oil palm plantations. RERTA is based in Riau Province, central Sumatra, Indonesia (0°56′0″ N, 101°18′0″ E), within industrial plantations owned and managed by PT Ivo Mas Tunggal (a subsidiary company of Golden Agri Resources, GAR – one of the largest oil palm companies globally). RERTA is replicated across two rivers, with treatments established during oil palm replanting (first-generation oil palms being replaced with young second-generation palms), first in 2018 and then in 2019 in the first and second river sites, respectively (Luke et al., 2020). At each river, we established three alternative restoration treatments and a control treatment, each covering an area of 400 m in length and 50 m in width on either side of the river. Treatments included: 1) enrichment plantings with a mix of six native tree species and remnant mature palms left in place in the margin (active restoration; hereafter Planting + MP), 2) mature palms left in place but no enrichment planting (passive restoration; hereafter MP only), 3) enrichment plantings with the same six native tree species, but mature palms removed (active restoration; hereafter Planting only), 4) control – mature palms removed and young oil palms replanted to the river’s edge (hereafter Replanted palms). Within the two enrichment planting treatments (Planting + MP & Planting only), an equal number of six native tree species (*Peronema canescens* (Sungkai); *Shorea leprosula (*Meranti tembaga); *Alstonia scholaris* (Pulai); *Artocarpus integer* (Cempedak); *Calophyllum inophyllum* (Bintangur); *Albizia chinensis* (Sengon)) were used, which were chosen from those that were readily available within regional nurseries, to provide a range of functional traits, including fruiting, leguminous, and pioneer species (Luke et al., 2020).

### 2.2. Field set-up

We established four 25 x 25 m sampling plots within each treatment in both replicate rivers. Two plots were positioned on either side of the river, each 12.5 m from the river’s edge. These plots were staggered from the north boundary of each treatment, running alongside the river at 50 m, 150 m, 250 m, and 350 m distances into each treatment (Figure 1).

**Figure 1:**
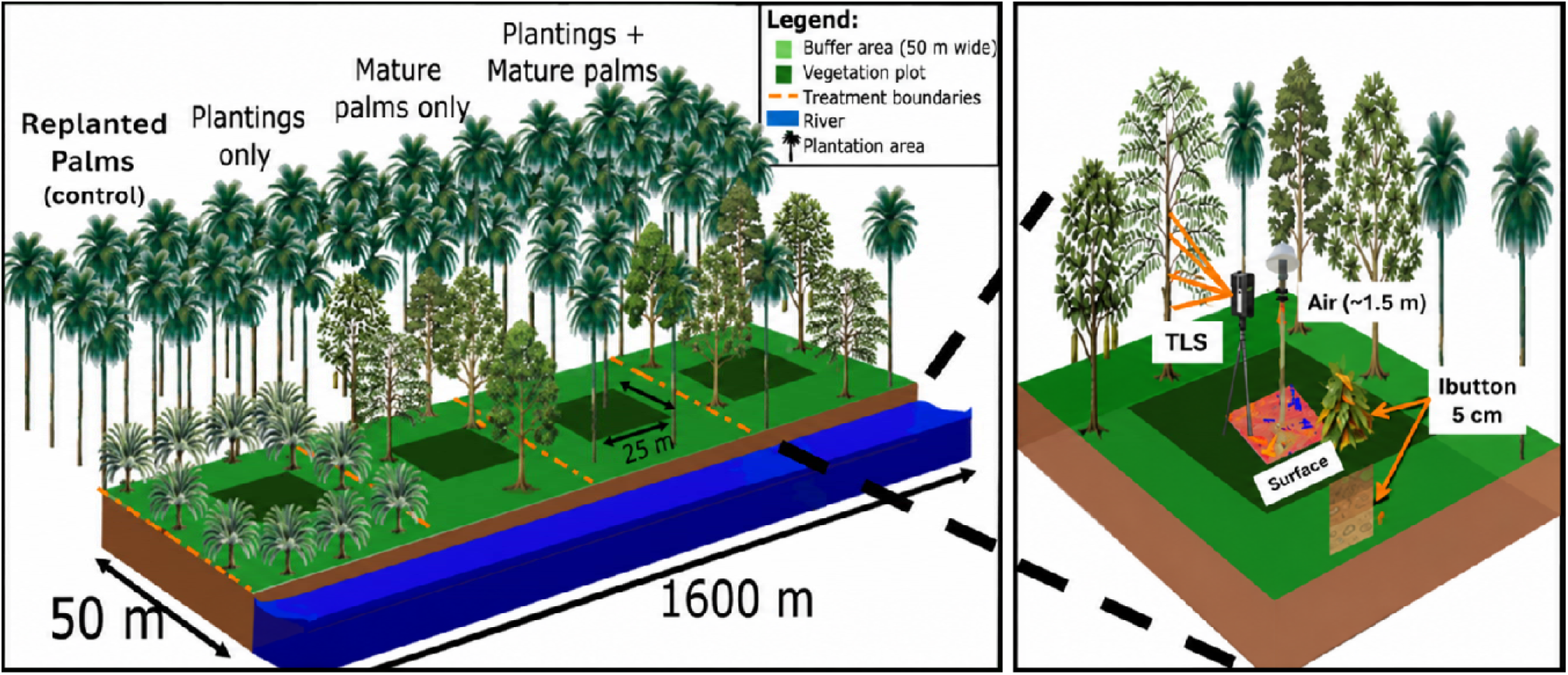
Summary schematic of the restoration treatments in the River Ecosystem Restoration in Tropical Agriculture Project (RERTA; left panel – example vegetation plots are not to scale) and an enlarged schematic of surveys completed within one of the four subplots per treatment (right panel). The RERTA Project includes three restoration treatments: Enrichment plantings + mature oil palms (Planting + MP), unassisted regenerating forestry with only remnant mature palms (MP only), assisted regeneration with enrichment plantings but the remnant mature palms removed (Planting only) and the control margin, with young oil palms planted to the river edge (Palms replanted). Enlarged plot surveys included vegetative complexity using a terrestrial LiDAR scanner (TLS) set at 1.5m above ground, temperature taken at three different heights (air, surface, and either in leaf litter or below-ground at 5cm depth).

#### 2.2.1. Vegetation structure across restoration treatments (Terrestrial Lidar Scanner)

To measure habitat structural characteristics, we used a FARO Focus 3D terrestrial laser scanner (Faro Technologies Inc., Lake Mary, USA), mounted on a tripod set at 1.3 m above ground in the centre of each sampling plot. The device was set to scan a field of view of 360 degrees horizontally and 305 degrees vertically, with an angular step width of 0.035, resulting in ∼44.4 million measurements per scan. Across the four treatments and two replicate river sites, we carried out 32 individual scans during the first field season (Dry Season = July 2023: one per sampling plot).

#### 2.2.2. Microclimate and land-surface temperature responses to restoration (Tmax, Tmean, Tmin, VPD, LST)

##### Air, leaf litter and below-ground temperature

Macroclimate and microclimate temperatures were split into specific height categories following Senior et al. (2018). We installed temperature loggers at three heights (1.5 m above ground, at ground level, and 5 cm below-ground) at the centre of each sampling plot, allowing us to assess and compare the impacts of alternative restoration treatments on different components of the ecosystem. Temperatures were recorded hourly for 14 consecutive days during the dry season (July 2023) and again during the wet season (February 2024). We were limited in the number of temperature-recording devices that available to deploy in the field, so we dispersed the devices to ensure approximately equal numbers of model types.

For air recordings, we used 13 HOBO temperature loggers (model: UA-001-08 8K; accuracy: 0.5°C, resolution: 0.1°C) and 20 humidity and temperature HOBO dataloggers (U23-001 model; temperature accuracy: 0.21; temperature resolution: 0.02; relative humidity accuracy: 2.5%; relative humidity resolution: 0.05), attached to bamboo stakes. To prevent exposure to direct solar radiation, temperature loggers were secured inside an overturned white plastic bowl, and humidity loggers were placed between two flat white plastic plates to allow horizontal airflow.

Directly beneath each HOBO temperature logger, we installed two iButton dataloggers (DS1921G-F5 Thermochrons) to measure ground-level temperatures: one placed just below the leaf litter layer (“ground level”) and another inserted into the below-ground at ∼5 cm depth. To account for systematic differences between logger models, we co-located additional iButtons at the same locations as a subset of randomly selected HOBO loggers. We used these paired measurements to calculate correction factors (see Section 2.3 for details). Again, we were limited in the number of iButtons available, so we aimed to distribute them evenly across treatments for leaf litter. Forty-eight iButtons were used, 16 in leaf litter (2 per treatment) and 32 in below-ground (4 per treatment). For each height level of dataloggers (air, leaf litter and soil), we placed iButtons in the centre of each of the four sampling plots (∼25 m from the river edge and spaced 50 m apart, running parallel to the river).

##### Surface temperature

We measured the surface temperature at the centre of each sampling plot using an infrared camera (FLIR E8-XT Systems). Each river site was visited over 2 days (4 days in total). We photographed each sampling plot twice a day in the morning (08:00 – 11:00) and afternoon (12:00 – 15:00). For each photograph, we stood at the centre of the plot, took a single hygrometer reading and then, holding the camera at breast height and pointing 45◦ downwards (relative to the ground), took a photo in four directions (90◦ turn each time). Each thermal image comprised 76,800 distinct observations of surface temperature (one per pixel) and covered a surface area of approximately 1 m^2^ (four photos combined = 307,200 observations). We recorded 256 thermal images (4 images per plot, 4 sampling plots per treatment, 4 treatments, 2 replicate rivers, and 2 time periods). We used Thermstats R package to calculate stats for each thermal image and then aggregated across the 4 images (Senior et al., 2019b). The temperature of cool and hot surface microclimates was defined as the lowest and highest 5^th^ percentile across all pixels.

##### Thermal buffering by treatments

Weather stations (DAVIS) were installed during the project setup in 2018 and 2019, along roadsides outside the riparian margins at each river site, thereby exposing them to direct solar radiation. By comparing data collected from these stations with data from Hobo and iButton loggers, we quantified the thermal buffering effect of riparian margins across restoration protocols.

### 2.3 Data Processing and Statistical Analyses

All analyses were performed in R (R Core Team, 2024) within RStudio. Data handling relied on tidyverse; mixed-effects modelling used lme4 with post-hoc contrasts from emmeans (Bates et al., 2015); figures were generated with ggplot2 (Wickham, 2016); spatial statistics for thermal imagery used spdep. Unless otherwise stated, tests were two-sided with α = 0.05.

#### 2.3.1 Vegetation structure across restoration treatments

Terrestrial laser scans were imported to FARO SCENE and registered using the standard algorithm. Registered point clouds were exported in XYZ format and processed in R to derive the Stand Structural Complexity Index (SSCI: See glossary in Supplementary for further details), Effective Number of Layers (ENL), canopy openness (CO), mean fractal dimension (MeanFrac), and canopy height (CH), following Ehbrecht et al. (2016). Treatment effects on SSCI, MeanFrac, and CO were analysed with generalised linear mixed-effects models, including Treatment as a fixed effect and River site as a random intercept. Because CO is a proportion, models used a binomial error with a logit link; overdispersion, where present, was addressed via an observation-level random effect. ENL and CH violated Gaussian assumptions and were therefore tested with Kruskal–Wallis tests followed, when significant, by Dunn’s post hoc comparisons. For mixed models, Tukey-adjusted pairwise contrasts were estimated with emmeans.

#### 2.3.2 Microclimate and land-surface temperature responses to restoration (Tmax, Tmean, Tmin, VPD, LST)

For each logger, we computed daily maxima, means, and minima for temperature and relative humidity (RH). To control for systematic differences among logger models, six iButtons were co-located with randomly selected HOBO loggers (U23-001: accuracy ± 2.5%, resolution 0.05%). Linear regressions of iButton on HOBO air temperatures were fitted for each season (dry: standard error = 2.107, *p* < 0.005; wet: standard error = 0.4093, *p* < 0.005). Below-ground and leaf-litter temperatures recorded by iButtons were then adjusted to the HOBO scale using the fitted intercept and slope (Adjusted T = Intercept + Slope × Original T), and all subsequent analyses for these strata use adjusted values (Supplementary S1).

Vapour pressure deficit (VPD) was calculated for each hourly air temperature/RH pair, as the difference between the saturation vapour pressure and the actual vapour pressure (Jucker et al., 2018; Montejo-Kovacevich et al., 2020). VPD was selected because it provides a direct measure of the atmospheric drying power, which is increasingly recognised as an important component of climate change and a key determinant of insect physiology, water balance, behaviour and thermal stress (Newell et al., 2023). Unlike relative humidity, VPD depends solely on temperature and humidity, simplifying interpretation and can be expressed as RH = (*e*/*e_s_*). *e_s_*. was calculated from temperature (in ◦C) using Bolton’s equation (Bolton, 1980).

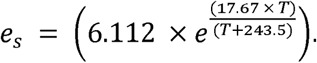

Temperature (daily max/mean/min) and VPD responses were analysed with linear mixed-effects models fitted separately for each height stratum, with Treatment and Date as fixed effects and Sub-plot ID (sampling plot) as a random intercept: Response ∼ Treatment + Date + (1|Sub-plot). A River Replicate random effect was excluded when it caused singular fits and explained negligible variance. Model assumptions were visually checked; the date was mean-centred or scaled as needed for convergence. Where overall Treatment effects were significant, Tukey-adjusted pairwise comparisons were reported.

##### Diurnal temperature range

For each logger and day, the diurnal temperature range (DTR) was calculated as the daily maximum temperature minus the daily minimum temperature, and then averaged across the recording period. DTR was analysed with the same model structure used for temperature and VPD (Response ∼ Treatment + Date + (1|Sub-plot)).

##### Thermal imagery: land-surface temperature and patch metrics

Surface thermal heterogeneity was quantified from thermal images by first computing the daily surface temperature range within each plot and then identifying local hot and cool patches using the Getis–Ord local Gi∗G_i^*Gi∗ statistic with queen’s contiguity (eight neighbours) implemented in spdep: localG (Bivand & Wong, 2018; Senior et al., 2019a). From these maps, we derived the patch temperature range (median of warmest minus median of coolest patches), mean cool-patch area (number of cool pixels × pixel area, averaged across four images), and an Aggregation Index describing the degree of clustering of cool patches, where higher values indicate stronger clustering in space (calculated as the proportion of shared edges among cool patches relative to the maximum possible; Senior et al., 2019b, 2019a). Patch metrics were analysed using mixed models that included Treatment, Time of day (morning/afternoon), and their interaction as fixed effects, with random intercepts for Site and Sub-plot.

##### Integrative multivariate analysis

To summarise vegetation structure, we conducted a principal component analysis on z-scored CH, CO, ENL, MeanFrac, and SSCI. PC1 (explaining 50.17% of the total variance) was used as a structural covariate to model daily maximum, mean, and minimum temperatures (analysed separately by season) using linear models that included Treatment, PC1, and their interaction. Our PCA of vegetative structure (Figure 6. A) showed PC1 was strongly positively associated with canopy height (r = 0.92), vertical structural complexity (ENL; r = 0.85), and stand structural complexity (SSCI; r = 0.60), and strongly negatively associated with canopy openness (r = −0.73). MeanFrac showed only a weak association with PC1 (r = −0.19) and contributed minimally to this axis. Non-significant interactions were removed, and assumptions were assessed via Shapiro–Wilk tests for residual normality and Levene’s tests for homogeneity of variances. All models met the assumptions of linear regression, with residuals showing no significant departure from normality (*p* > 0.36 for all Shapiro-Wilk tests) and equal variance among treatments (*p* > 0.14 for all Levene’s tests).

#### 2.3.3 Thermal buffering by treatments: present offsets and future projections relative to insect CTmax

##### Present-day temperature offsets

To quantify microclimatic buffering, we calculated daily temperature offsets between *in situ* loggers within riparian treatments and a nearby open-plantation weather station (DAVIS) as margin minus outside temperature for daily maxima, means, and minima (Offset = Tmargin \ mathrm {margin} margin – Toutside \mathrm {outside} outside). To project buffering under future climates, we combined observed offsets with downscaled CHELSA v2.1 temperatures retrieved via pastclim (Leonardi et al., 2023). Station data from 2019–2024 (51 months) were validated against historical CHELSA (1980–2010) at the field site (101.43° E, 0.93333° N), yielding a mean annual difference of 0.4 °C; this delta was added to all CHELSA temperatures used in projection (Supplementary S2). We extracted temperature projections at 0.5′ resolution (∼1 km) for 2045 and 2085 under SSP1-2.6, SSP3-7.0, and SSP5-8.5 using the UKESM1-0-LL global climate model. Future buffered temperatures were estimated as T_buffered,future_ = T_ambient,future_ + Offset_observed_; we then computed ΔBuffer = T_buffered,future_ – T_ambient,future_ for comparisons among treatments and time horizons.

##### Future temperature offsets

To assess biological relevance, we compared projected buffered temperatures against published tropical insect thermal limits. Critical thermal maxima (CTmax) and thermal optima (Topt) were compiled using existing published data extracted from GlobTherm (Bennett et al., 2018) and Deutsch et al. (2008) for multiple insect orders (Diptera, Hymenoptera, Odanata, Hemiptera) and knockdown assays of tropical butterflies ((Ashe-Jepson et al., 2023; see Supplementary S4 for details of publications and data used). These comparisons are based on published limits across multiple species and regions, grouped by order, rather than knockdown assays we conducted as part of this project. Therefore, these predicted limits should be interpreted as risk indicators rather than direct outcomes within our river forest experimentation or from the study region specifically. We quantified proximity to thermal limits for late-century SSP5-8.5 as Proximity_CTmax_ = CTmax − T_buffered,max_. We did not repeat the statistical comparison between treatments, as our method for calculating future climate conditions would maintain the same differences as modern climate conditions.

## 3. Results

### 3.1. Vegetation structure across restoration treatments

All measured metrics of forest structural complexity, stand density, effective number of layers, MeanFrac, CO, and CH, varied significantly between alternative restoration treatments (Fig. 2a–e). Stand density differed significantly by treatment (ANOVA: χ*²* = 65.71, *df* = 3, *p <* 0.001; Fig. 2a), with the highest densities observed in actively restored margins (Planting + MP higher than palms removed). These values were significantly greater than those in passive restoration (MP only) and control (replanted palms) treatments (GLM post hoc: β = 0.10, *SE* = 0.01, *df* = 27, *p <* 0.001). ENL also varied across treatments (Kruskal–Wallis: χ*²* = 17.49, *df* = 3, *p <* 0.001; Fig. 2b), with greater vertical layering in actively restored margins than in the control. MeanFrac showed no significant difference between the two active restoration treatments (ANOVA: χ*²* = 32.23, *df* = 3, *p =* 0.78), but both differed significantly from the passive and control treatments (ANOVA: χ*²* = 32.23, *df* = 3, *p <* 0.001), with both being intermediate to the passive and control treatments (Fig. 2c). Canopy openness varied significantly across treatments (GLM: χ*²* = 21.10, *df* = 3, *p <* 0.0001; Fig. 2d), increasing in order from Planting + MP, Planting only, MP only, and Replanted palms. Canopy height was significantly greater across all restoration treatments than in the control (Kruskal–Wallis: χ*²* = 18.56, *df* = 3, *p <* 0.005; Fig. 2e).

**Figure 2:**
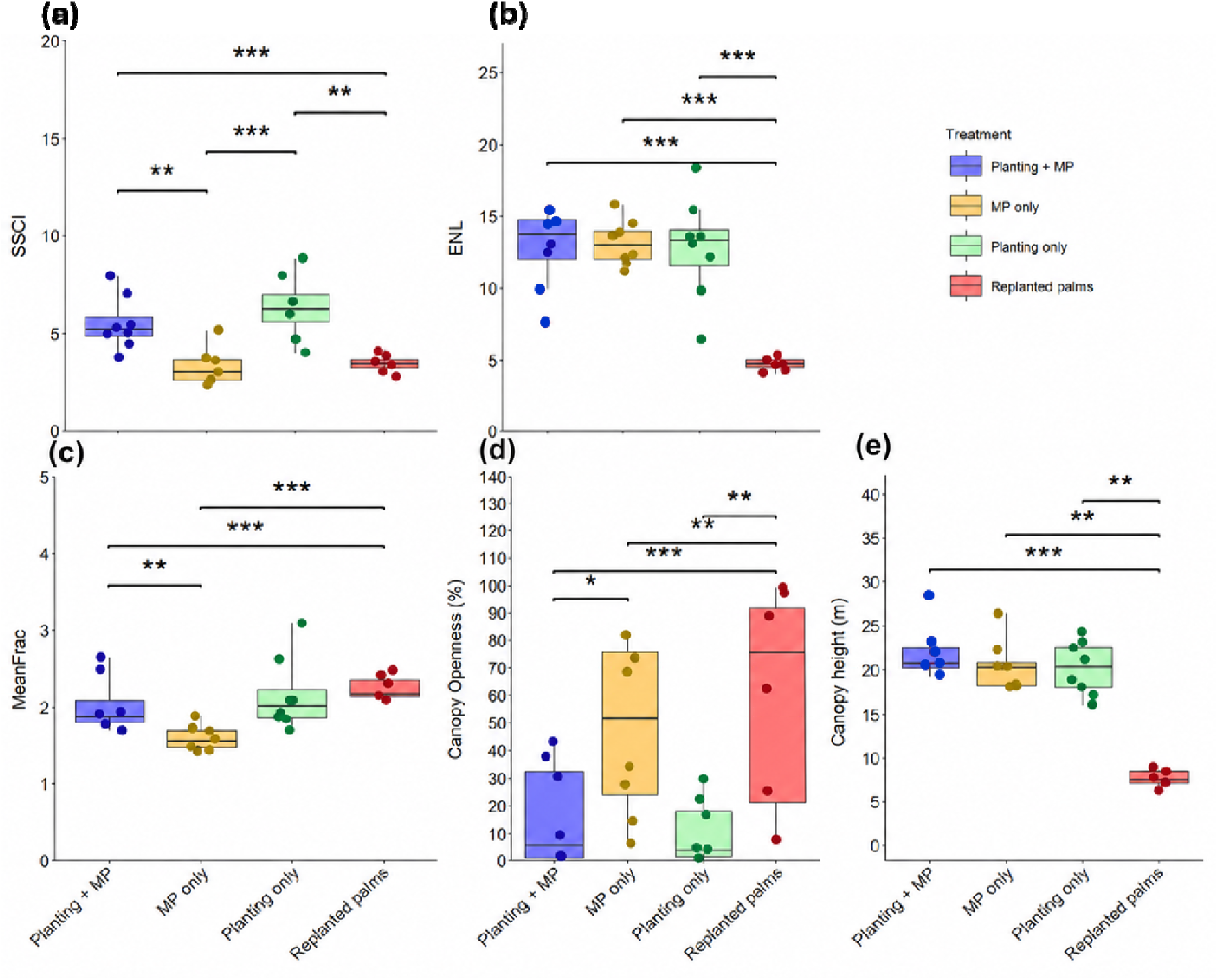
Vegetative complexity metrics across restoration treatments. **(a)** Stand Structural Complexity Index (SSCI), **(b)** Effective Number of Layers (ENL), **(c)** Mean Fractal Dimension (MeanFrac), **(d)** Canopy Openness, and **(e)** Canopy Height. Whiskers extend to the minimum and maximum values within 1.5 times the IQR. Points beyond this range are plotted as individual outliers. Significant differences are denoted by asterisks (*; Tukey *post hoc* test, *p <* 0.05). Shorthand expressions indicate the treatment pair being compared, with asterisks marking the level of significance (\*\*\**p <* 0.001, ** *p <* 0.01,* *p <* 0.05*) and †* indicating non-significance. Metrics were calculated for each restoration treatment: Planting + MP: mature oil palm with enrichment planting; MP only: mature oil palm only; Planting only: mature oil palm removed with enrichment planting; Replanted Palms (control): replanted oil palms planted to the river edge) and averaged across sites and sampling plots.

### 3.2. Microclimate and land-surface temperature responses to restoration (Tmax, Tmean, Tmin, VPD, LST)

#### 3.2.1. Air temperature (1.5 m aboveground)

In the dry field season, daily maximum and mean ambient air temperatures varied significantly across restoration treatments (Fig. 3B–C), while minimum temperatures did not (LMM: χ*²* = 7.57, *df* = 3, *p =* 0.056; Fig. 3D). Maximum air temperature was significantly lower in actively restored margins with enrichment plantings (Planting + MP and Planting only) than in both passive restoration (MP only) and control margins (replanted palms) (LMM: χ*²* = 15.05, *df* = 3, *p =* 0.0018; Fig. 3B). Mean air temperature was also significantly lower in both actively restored margins and the passive margin treatment relative to the control, with replanted palms exhibiting the highest mean temperatures (LMM: χ*²* = 18.80, *df* = 3, *p =* 0.001; Fig. 3C).

**Figure 3:**
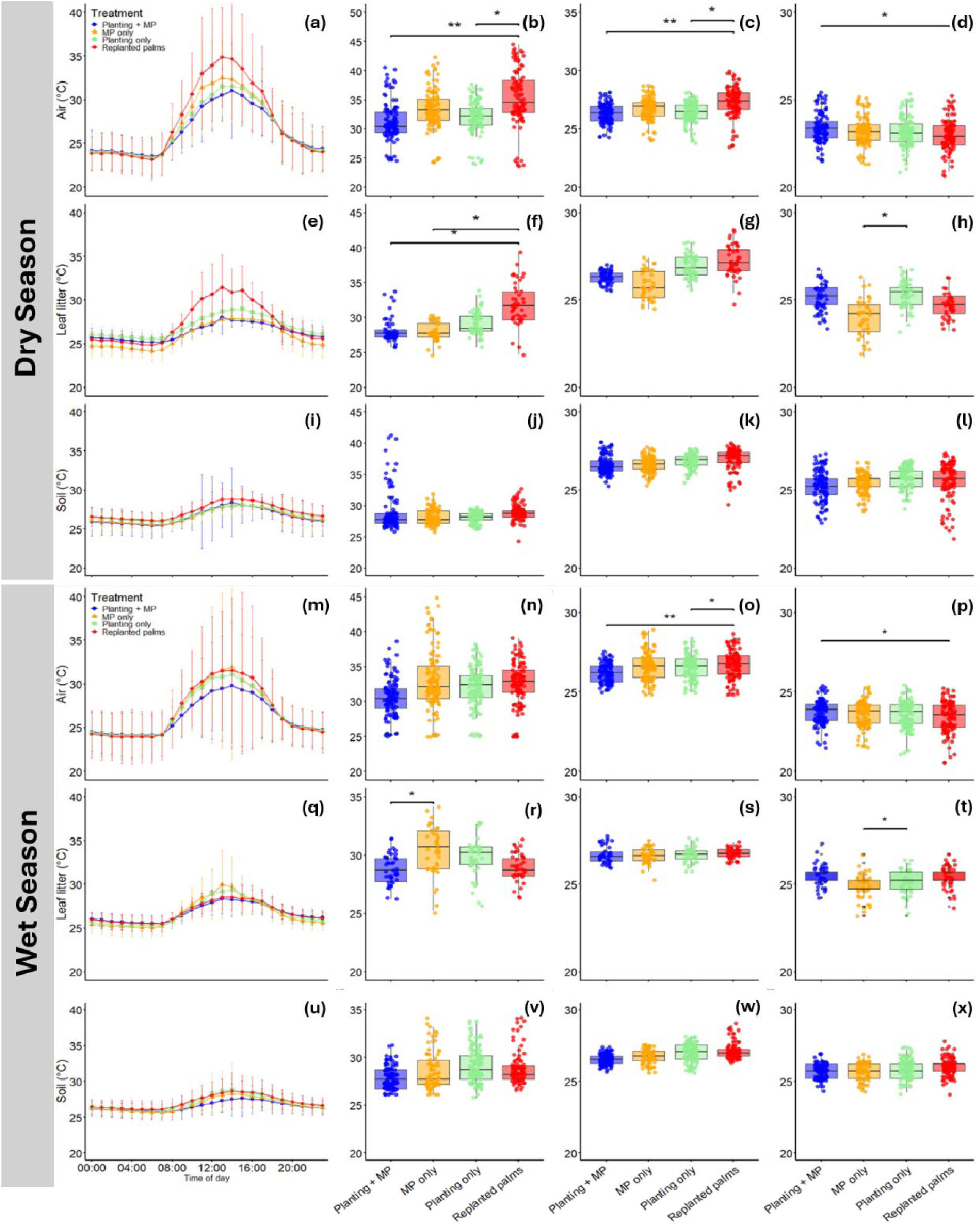
Daily temperatures across restoration treatments and field seasons. Hourly temperature variation was recorded every hour for air **(a, m)**, leaf litter **(e, q)**, and below-ground **(i, u)**. The mean daily maximum, mean, and minimum temperature from the two field seasons, dry (top panel) and wet (bottom panel), across the restoration treatments: Planting + MP: mature oil palm with enrichment planting; MP only: mature oil palm only; Planting only: mature oil palm removed with enrichment planting; Palms replanted (Control): oil palms planted to river edge. For the hourly line plots, each coloured line represents a different treatment averaged across loggers placed in each sampling plot; the temperature was then averaged for each hour. Error bars were calculated by averaging the variance each hour or day and adding/subtracting from the points. For boxplots, the whiskers extend to the minimum and maximum values within 1.5 times the IQR. Points beyond this range are plotted as individual outliers. Data with significant statistical differences are shown by * above pairs (Tukey post hoc test, *p <* 0.05). Shorthand expressions indicate the treatment pair being compared, with asterisks marking the level of significance (\*\*\**p <* 0.001, ** *p <* 0.01,* *p <* 0.05).

#### 3.2.2. Leaf litter temperature (5 cm depth)

In the dry season, daily maximum and mean leaf litter temperatures differed significantly across restoration treatments (maximum: LMM, χ*²* = 18.62, *df* = 3, *p <* 0.001; mean: LMM, χ*²* = 13.75, *df* = 3, *p =* 0.003), with both metrics being lower in actively restored margins (Planting + MP and Planting only) than in passive restoration (MP only) and control sites (replanted palms; Fig. 3a). Minimum leaf litter temperature did not differ significantly among treatments (LMM: χ*²* = 7.57, *df* = 3, *p =* 0.056; Fig. 3g).

In the wet season, significant treatment effects were observed for maximum (LMM: χ*²* = 17.21, *df* = 3, *p <* 0.001), mean (LMM: χ*²* = 13.75, *df* = 3, *p =* 0.003), and minimum leaf litter temperatures (LMM: χ*²* = 17.10, *df* = 3, *p <* 0.001; Fig. 3b). Post hoc tests showed that passive restoration (MP only) had higher maximum and minimum leaf litter temperatures than the control treatment, while active treatments exhibited lower minimum temperatures than the control. No significant pairwise differences were detected for mean leaf litter temperature, despite a significant overall treatment effect. Full post hoc comparisons are provided in Supplementary Table S4.

#### 3.2.3 Below-ground Temperature (5cm depth)

No significant differences in maximum, mean, or minimum below-ground temperature were observed among treatments in either year (all p > 0.17; Fig. 3j–l, v–x; Supplementary Table S4).

#### 3.2.4 Diurnal Temperature Range

In the dry season, diurnal temperature range differed significantly among treatments for ambient air (ANOVA: χ*²* = 8.15, *df* = 3, *p =* 0.040) and leaf litter (ANOVA: χ*²* = 22.27, *df* = 3, *p <* 0.001), but not for below-ground (ANOVA: χ*²* = 1.87, *df* = 3, *p =* 0.60). The largest temperature ranges occurred in the control treatment (Replanted palms), followed by progressively smaller ranges in passive restoration (MP only), active restoration without mature palms (Planting only), and active restoration with mature palms (Planting + MP).

In the wet season, only leaf litter exhibited a significant treatment effect (ANOVA: χ*²* = 24.46, *df* = 3, *p <* 0.05), with the largest diurnal ranges again occurring in replanted palms and the smallest in Planting + MP. Diurnal temperature ranges for ambient air and below-ground were not significantly different among treatments (air: χ*²* = 7.66, *df*= 3, *p =* 0.053; below-ground: χ*²* = 4.39, *df* = 3, *p =* 0.22). Full post hoc results are detailed in Supplementary Table S6.

#### 3.2.5 VPD

Significant differences in mean and maximum daily VPD were observed across restoration treatments in both field seasons (mean: ANOVA, χ*²* = 22.50, *df* = 3, *p <* 0.001; maximum: χ*²* = 9.00, *df* = 3, *p =* 0.020; Fig. 4 a–d). In both dry and wet field seasons, the control treatment (Replanted palms) consistently exhibited higher mean and maximum VPD values than all restored treatments, while Planting + MP had the lowest VPD values. Full details on relative humidity differences by treatment and season are reported in Supplementary Table S5.

**Figure 4:**
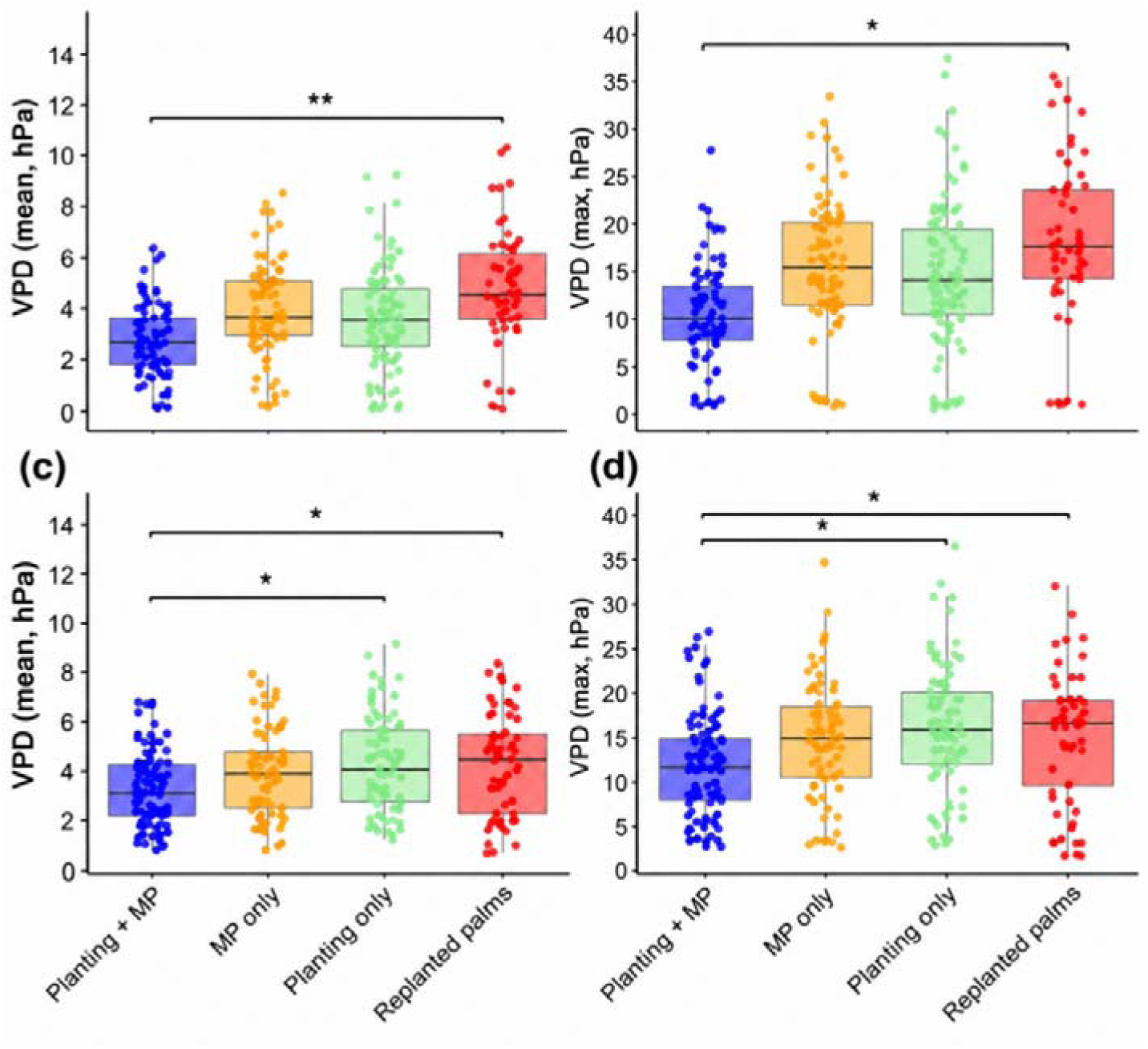
Vapour pressure deficit (VPD, ‘drying power’, hPa) across the microclimates sampled during the dry (top panel; a & b) and wet field seasons (bottom panel; c & d) across the restoration treatments: Planting + MP: mature oil palm with enrichment planting; MP only: mature oil palm only; Planting only: mature oil palm removed with enrichment planting; Replanted palms (Control): oil palms planted to river edge). Whiskers extend to the minimum and maximum values within 1.5 times the IQR. Points beyond this range are plotted as individual outliers. Statistically significant pairwise comparisons between treatments are denoted by * above the plots (Tukey *post hoc* test, *P <* 0.05). Shorthand expressions indicate the treatment pair being compared, with asterisks marking the level of significance (\*\*\**p <* 0.001, ** *p <* 0.01,* *p <* 0.05).

#### 3.2.6. Surface temperature Surface thermal characteristics

Median surface temperature varied significantly with treatment and time of day, with a significant interaction (LMM: χ*²* = 11.33, *p =* 0.010; Fig. 5a). Planting + MP consistently exhibited lower median surface temperatures than all other treatments at both times of day (all p ≤ 0.002). Temperature heterogeneity (SHDI) also differed by treatment and time of day, with a significant interaction (χ*²* = 14.94, *p =* 0.002; Fig. 5b); Planting + MP produced more uniform surface temperatures (lower SHDI) than all other treatments (*p <* 0.001).

**Figure 5:**
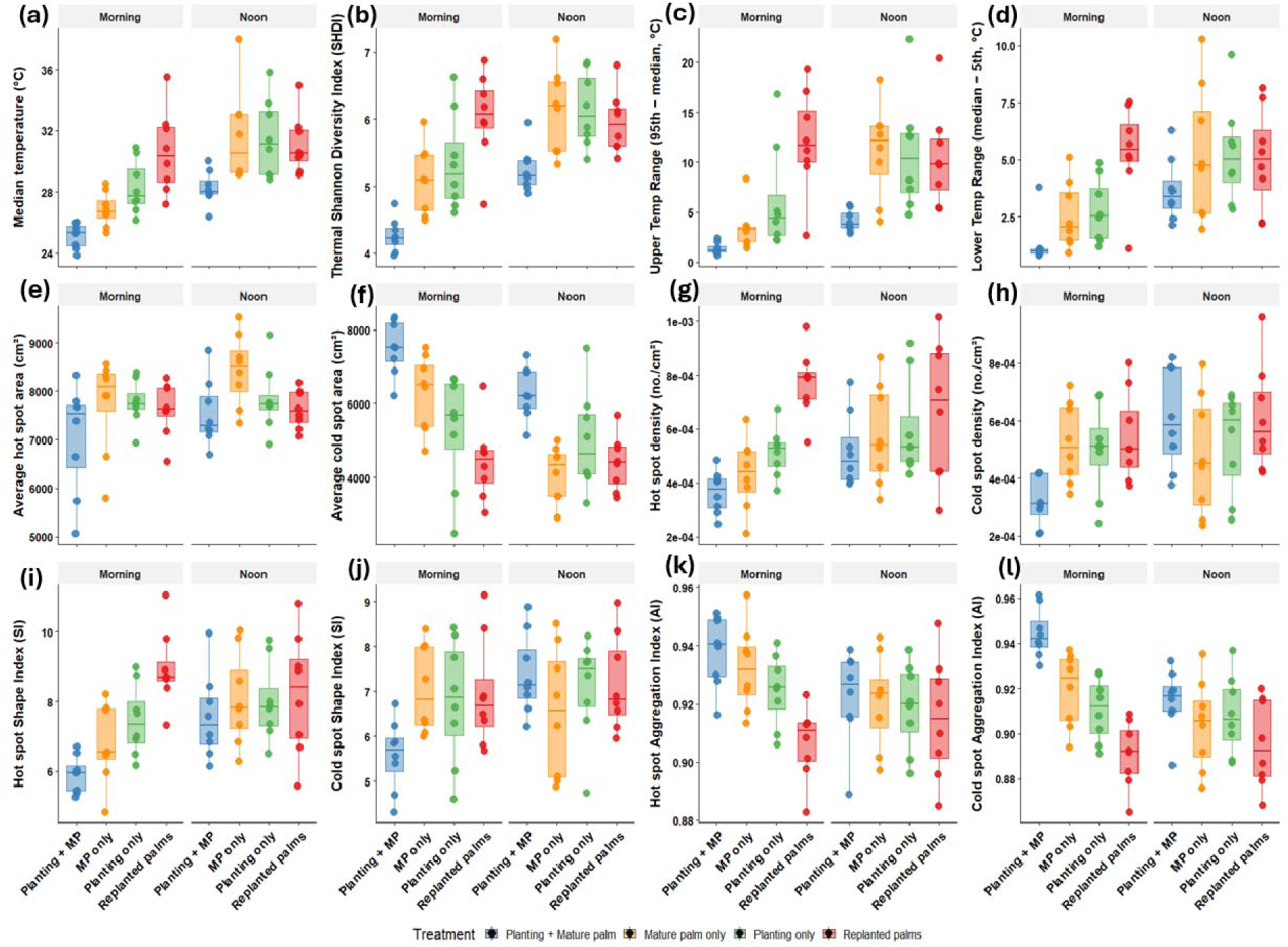
Trends in land surface temperature by the time of day and across treatments (Planting + MP (blue): enrichment planting with mature oil palm; MP only (yellow): mature oil palm only; Planting only (green): mature oil palm removed with enrichment planting; Replanted palms (Control; red): oil palms planted to river edge). Thermal camera was used to record characteristics at two points in the day (morning (8:00 – 11:00) and afternoon (12:00 – 15:00). From the left to right and top to bottom, the metrics are: median temperature **(a)**; thermal Shannon Diversity Index (SHDI) **(b)**; Upper Temp Range (95^th^ percentile minus median temperature) (c); Lower Temp Range (median temperature minus 5^th^ percentile) **(d)**; the average area (cm^2^) per hot spot **(e)**; the average area (cm^2^) per cold spot **(f)**; the number of hot spots per unit area **(g)**; the number of cold spots per unit area **(h)**; the Shape Index (SI) of hot spots **(i)**; the Shape Index (SI) of cold spots **(j)**; the Aggregation Index (AI) of hot spots **(k)**; the Aggregation Index (AI) of cold spots **(l)**. Whiskers extend to the minimum and maximum values within 1.5 times the IQR. Points beyond this range are plotted as individual outliers.

**Figure 6.**
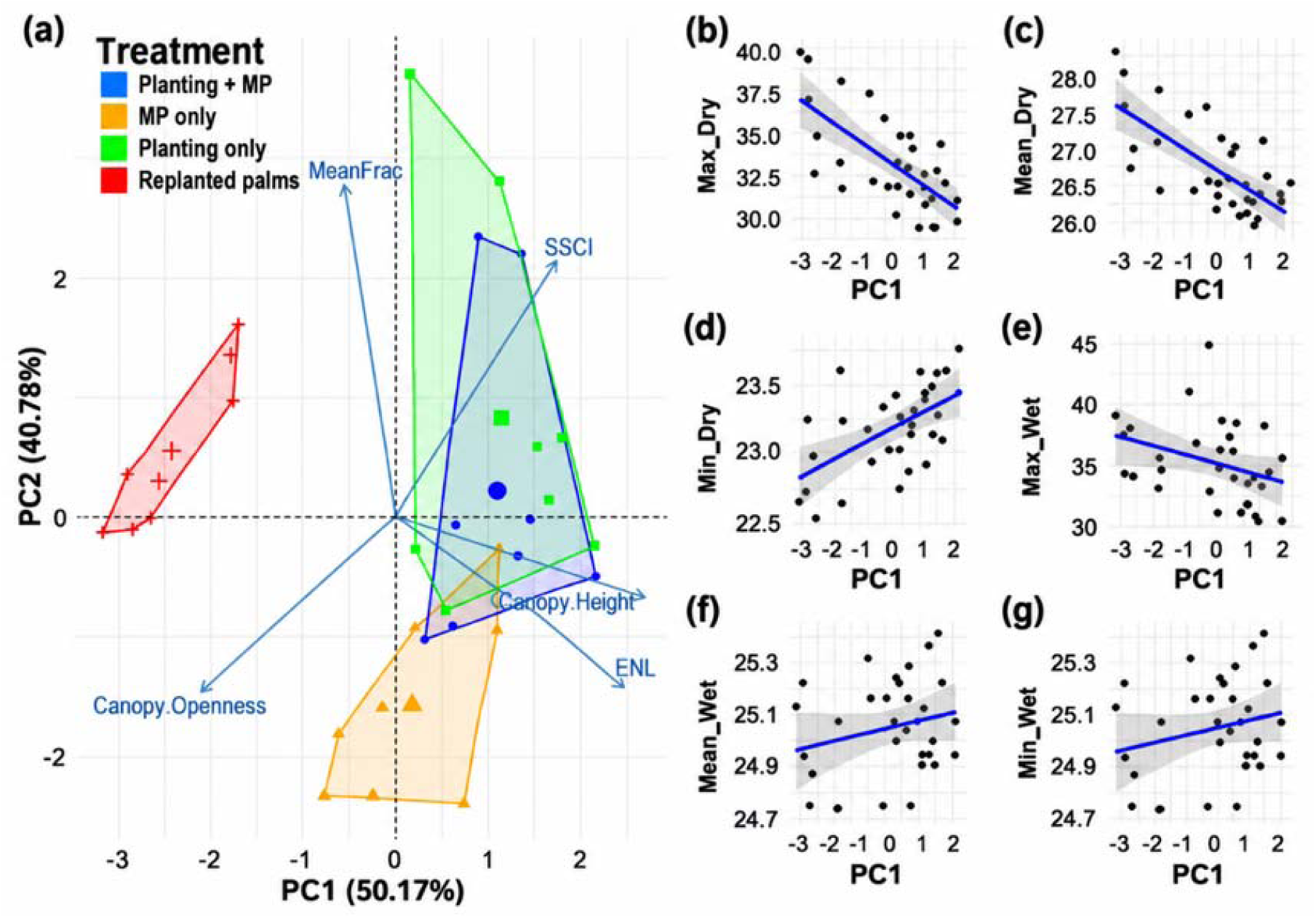
(a) Principal Component Analysis (PCA) of vegetative structural characteristics derived from terrestrial LiDAR scanning (n = 32), illustrating multivariate variation in vegetation structure grouped by restoration treatments (Planting + MP (blue): enrichment planting with mature oil palm; MP only (yellow): mature oil palm only; Planting only (green): mature oil palm removed with enrichment planting; Replanted palms (Control; red): oil palms replanted to river edge). PC1 reflects gradients in canopy openness, vertical complexity, and ground cover. Arrows represent the direction and strength of correlations between LiDAR-derived variables: Canopy Openness, Effective Number of Layers (ENL), MeanFrac, and Stand Structural Complexity Index (SSCI). Ellipses denote 95% confidence intervals around treatment centroids. **(b – g)** are linear regressions (n = 192) showing relationships between plot and macroclimate temperatures’ daily maximum and mean in the dry (Plot **b – d**) and wet season (Plot **e – g**).

##### Temperature variability

The upper temperature range showed significant effects of treatment, time of day, and their interaction, with Planting + MP reducing extremely high surface temperatures relative to all other treatments (all *p* ≤ 0.012; Fig. 5c). The lower temperature range also differed among treatments, being reduced in Planting + MP relative to replanting young palms (*p =* 0.001; Fig. 5d), although the treatment × time interaction was not significant.

##### Patch characteristics

Hot patch area differed among treatments (χ*²* = 11.44, *p =* 0.009; Fig. 5e), being smallest in Planting + MP, while cold patch area was also smallest in this treatment across both times of day (χ*²* = 12.09, *p =* 0.007; Fig. 5f). Hot patch density was lowest in Planting + MP (*p* ≤ 0.022; Fig. 5g), whereas cold patch density varied with time of day and its interaction with treatment, but did not differ significantly among treatments (Fig. 5h).

##### Patch shape and aggregation

Hot patch shape index differed among treatments, with Planting + MP producing simpler patch shapes than replanted palms (*p =* 0.0002; Fig. 5i). Cold patch shape index showed no apparent treatment effects (Fig. 5j). Hot patch aggregation index was lowest in Planting + MP compared to all other treatments (*p* ≤ 0.023; Fig. 5k), while cold patch aggregation index also varied by treatment, with the lowest values observed in Planting + MP (*p* ≤ 0.013; Fig. 5l).

#### 3.2.6 Cross analyses between stand structural complexity and temperature

Principal component analysis showed distinct clustering of vegetation structural complexity across restoration treatments (Figure 6a). The first principal component (PC1) was strongly positively correlated with canopy height (*r* = 0.92, *p* < 0.001) and the effective number of layers (ENL; *r* = 0.85, *p* < 0.001), and negatively correlated with canopy openness (*r* = −0.73, *p* < 0.001), indicating that increasing PC1 scores represented increasingly structurally complex and closed-canopy vegetation (Supplementary Table S8). However, no significant interaction between vegetation structural complexity (PC1) and restoration treatment was detected for any temperature metric (all *p* > 0.12; Supplementary Table S6). Significant positive associations were observed between PC1 and maximum temperature in the dry season (*F*(1,28) = 22.59, *p <* 0.001), mean temperature (*F*(1,28) = 22.92, *p <* 0.001), and maximum temperature in the wet season (*F*(1,28) = 5.10, *p =* 0.031). Minimum temperature in the dry season was also positively associated with PC1 (*F*(1,28) = 13.12, *p <* 0.001, *R*² = 0.32). In contrast, mean and minimum temperatures in the following wet season showed no significant relationships with vegetation structure (both *F*(1,28) = 2.14, *p =* 0.156).

#### 3.2.7 Thermal buffering by treatments: present offsets and future projections relative to insect CTmax

##### 3.2.7.1. Modern climate buffering

Daily temperature extremes differed significantly among restoration treatments. Daily maximum buffered temperatures varied by treatment (LMM: *F*(3,12.1) = 7.84, *p =* 0.0036), with planting with mature palms exhibiting lower daily maxima than Mature Palms Only (–4.77 °C, SE = 1.33, *p =* 0.012) and replanting young palms (–4.49 °C, SE = 1.33, *p =* 0.018).

Daily mean buffered temperatures also differed among treatments (LMM: *F*(3,12.5) = 8.13, *p =* 0.0029), with planting with mature palms being cooler than replanting young palms (–0.80 °C, SE = 0.19, *p =* 0.0032), while Mature Palms Only and Planting Only were intermediate and did not differ significantly from either extreme.

Daily minimum buffered temperatures showed the strongest treatment effect (LMM: F(3,13) = 19.88, *p <* 0.0001). Planting with mature palms maintained warmer nighttime conditions than Planting Only (0.23 °C, SE = 0.07, *p =* 0.023) and replanting young palms (0.46 °C, SE = 0.07, *p <* 0.0001). Mature Palms Only also retained more nighttime heat than replanting young palms (0.32 °C, SE = 0.07, *p =* 0.0013).

For maximum temperature offsets (Fig. 7), outside temperature and treatment showed a strong interaction (χ*²* = 280.47, *df* = 3, *p <* 0.001), alongside significant effects of outside temperature (χ*²* = 482.98, *df* = 1, *p <* 0.001) and treatment (χ*²* = 160.13, *df* = 3, *p <* 0.001). Mature Palms Only and replanting young palms exhibited larger positive maximum temperature offsets than planting with mature palms (both *p <* 0.001), indicating reduced thermal buffering. Mean temperature offsets showed a significant interaction (χ*²* = 16.89, *df* = 3, *p =* 0.001) and treatment effect (χ*²* = 163.72, *df* = 3, *p <* 0.001) but not of outside temperature (χ*²* = 0.51, *df* = 1, *p =* 0.476); Mature Palms Only and replanting young palms again exhibited larger mean temperature offsets than planting with mature palms (both *p <* 0.001). Minimum temperature offsets were influenced by outside temperature (χ*²* = 67.72, *df* = 1, *p <* 0.001) and treatment (χ*²* = 57.16, *df* = 3, *p <* 0.001), with no interaction (χ*²* = 3.99, *df* = 3, *p =* 0.263). Planting with mature palms maintained higher minimum temperature offsets than both Planting Only and replanting young palms (both *p <* 0.01).

**Figure 7.**
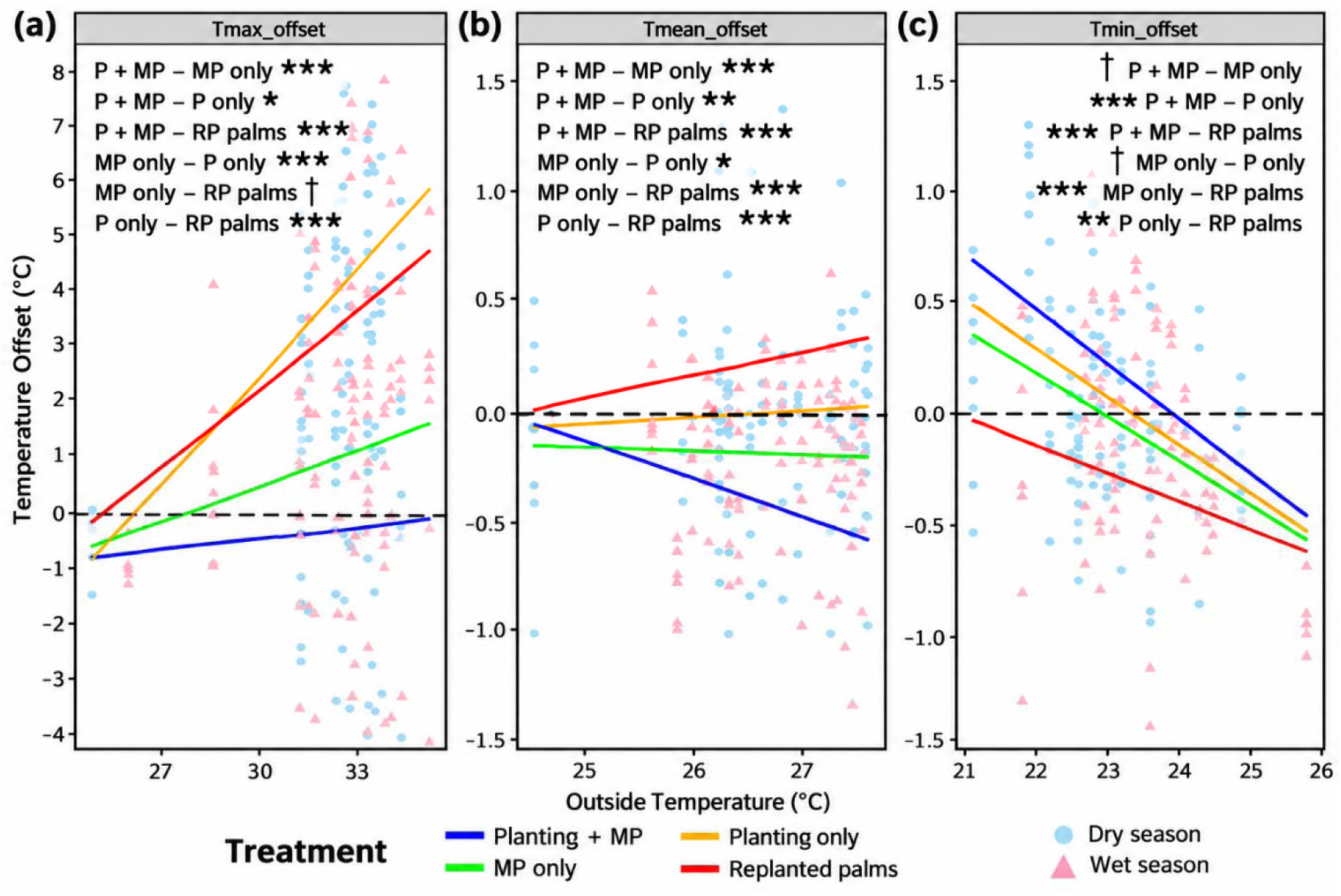
Thermal offsets in the margin compared to the outside environment. Panels show daily maximum **(a)**, mean **(b)**, and minimum **(c)** temperature offsets (temperature difference between inside and outside the temperature range: y-axis) across restoration treatments compared to the temperature outside the margin (x-axis). Data are grouped by year, with the dry season represented by circles (●) and wet season by triangles (▲). Restoration treatments are distinguished by line colour: Planting + MP (blue); enrichment plantings with mature oil palms, MP only (green); mature oil palms only, Planting only (orange); old oil palms removed and replanted with seedlings, and Replanted palms (Control; red): oil palms replanted. **Significance annotations above each panel denote pairwise comparisons between treatments; shorthand expressions indicate the treatment pair being compared, with asterisks marking the level of significance (\*\*\**P <* 0.001, \**P <* 0.01*, P <* 0.05*) and †* indicating non-significance. Letters denote overall differences among treatments based on Tukey post hoc tests (*p <* 0.05).

##### 3.2.7.2. Future climate thermal buffering

Across all scenarios and both mid– and late-century projections, planting with mature palms produced the greatest positive thermal buffering (+0.41 °C), followed by Planting only (+0.17 °C), MP only (–0.04 °C), and Replanted palms (–0.39 °C), which consistently exhibited the weakest buffering under absolute air temperature conditions. Consistent with these differences, thermal safety margins were narrowest in Replanted palms (Fig. 8). Mean body temperatures in this treatment were 1.35 °C closer to CTmax than in planting with mature palms, 1.22 °C closer than in MP only, and 1.36 °C closer than in Planting only.

**Figure 8:**
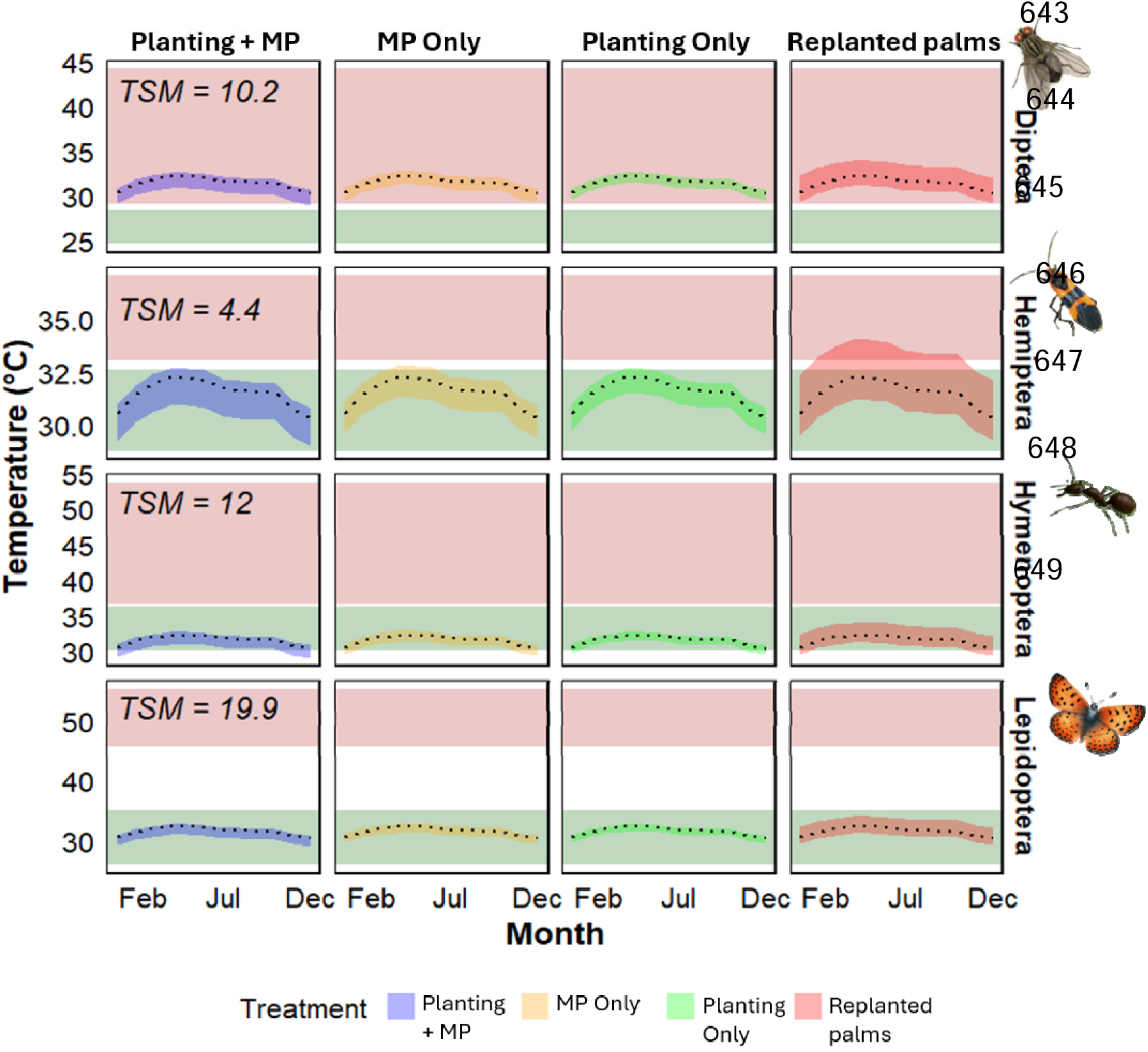
Future projected microclimate temperatures (°C) across insect orders and restoration treatments (Planting + MP; mature oil palm with enrichment planting, MP Only; mature oil palm only, Planting Only; mature oil palm removed with enrichment planting, and Replanted palms (Control; young oil palms planted to the river edge)). Projections are based on the UK GCM under the high-emissions SSP585 scenario for the late-century period (∼2085). Red shaded bands represent the critical thermal maximum range (CTmax) for each insect order based on published data (Supplementary S4); green bands represent their estimated thermal optimum (Topt). The black dotted line indicates macroclimatic station temperature. Shaded ribbons show predicted buffered microclimate temperatures inside riparian restoration plots. Thermal safety margins (TSM: distance between environmental/body temperature and thermal limit) are shown in the top-left panel of each insect order.

## 4. Discussion

Across the first 5–6 years of restoration, riparian margins diverged markedly in both vegetative structure and thermal regimes. Active restoration accelerated canopy development and vertical layering, and these structural differences were consistently associated with cooler and more stable microclimates. Effects were strongest where enrichment planting was combined with retained mature palms (Planting + MP), followed by Planting only; passive restoration (MP only) was intermediate, and the replanted young palms (control) had the lowest structural complexity and most variable microclimate. Canopy openness emerged as the dominant structural correlate of temperature across seasons, highlighting canopy closure as a key variable in climate-buffering effectiveness to manage for in working tropical landscapes.

### 4.2 Drivers of riparian forest characteristics

Active restoration treatments exhibited higher canopies, greater vertical stratification (ENL), and lower openness than passive and control treatments, consistent with evidence that enrichment planting accelerates the development of multilayered forest structure and associated ecosystem functions (Crouzeilles et al., 2020; Holl & Brancalion, 2020; Meli et al., 2017). Structural gains translated into lower maximum and mean air and litter temperatures, reduced vapour-pressure deficit (VPD), and narrower diurnal temperature ranges, particularly during hours of peak temperature (early afternoon; 12 – 15:00). These outcomes align with well-established mechanisms whereby closed, multilayered canopies intercept shortwave radiation, dampen convective heating, and enhance evaporative cooling via transpiration (De Frenne et al., 2019; Jucker et al., 2018)). In contrast, below-ground temperatures did not differ among treatments, likely reflecting thermal inertia (higher heat capacity and attenuated diurnal amplitude) and the shallow soil layer’s weaker direct sensitivity to canopy shading compared to the aboveground strata.

Overall our results revealed that Planting + MP and Planting only were associated with higher canopy height, greater layering, and lower openness, and these structural gradients were predictably linked to cooler maximum and mean temperatures. Notably, we detected no interaction between treatment and the structure–temperature relationship, indicating that treatments act primarily by repositioning plots along a common structural pathway, rather than altering this relationship (De Frenne et al., 2019; Jucker et al., 2018; Senior et al., 2017). Night-time minima were less consistently linked to structure, suggesting a larger role of landscape topography, wind, or soil moisture in governing nocturnal cooling.

### 4.3 Surface Temperature

Our results demonstrate a strong association between restoration treatments and both the magnitude and spatial distribution of surface temperatures. Thermal imagery showed that Planting + MP produced fewer and smaller hot patches, alongside more aggregated and extensive cool patches, relative to other treatments. These patterns indicate not only lower mean surface temperatures but also the presence of denser cool microrefugia at fine spatial scales, supported by increased vegetation structure, moisture retention, and vertical layering. Not only are cool microrefugia important, but animals that require hotter conditions can also access suitable conditions at shorter distances within ecosystems that provide more varied thermal conditions (Scheffers et al., 2017).

Surface temperature dynamics were analysed in parallel with ambient air temperature because insects are susceptible to surface thermal conditions and can utilise these to thermoregulate (Kerr et al., 2025). As small-bodied ectotherms, insects experience rapid heat gain from substrates and are particularly vulnerable to surface temperature changes, owing their reliance on flight for movement, combined with high metabolic demands. Consequently, surface thermal conditions may impose stronger constraints on insect activity and habitat use than ambient air temperature alone (Leclerc et al., 2022; Scheffers et al., 2017).

Beyond differences in absolute temperatures, restoration treatments also altered surface thermal heterogeneity. Under standard oil palm conditions, the thermal landscape may restrict habitat suitability for taxa sharing the same spatial domain, such as along river buffers, by limiting the availability of accessible thermal refuges within short movement distances. This indicates that fine-scale surface thermal conditions could constitute an important component of habitat quality for invertebrates, and should be considered in management policies in addition to traditional measures of habitat availability (Scheffers et al., 2017).

### 4.4 Relevance under climate change: projections and insect thermal risk

Using observed microclimate offsets relative to an open-plantation station and downscaled CHELSA projections, we found that restored margins, particularly Planting + MP, consistently provided the strongest thermal buffering under mid– and late-century climate scenarios, including SSP5-8.5. To assess biological relevance, projected buffered temperatures were evaluated against published thermal limits (CTmax and Topt) for tropical insects (Ashe-Jepson et al., 2023; Bennett et al., 2018; Deutsch et al., 2008). Across treatments, replanted young palms exhibited the narrowest thermal safety margins for a range of insect orders, whereas margins with mature palms were the farthest from critical thermal thresholds.

Projected exposure to thermal stress varied among insect orders, reflecting differences in thermal tolerance and adaptive strategies. Our modelling suggests that Diptera and Hemiptera are likely to experience elevated thermal risk in established oil palm plantations under future climate conditions. At the same time, Hymenoptera and Lepidoptera generally remain within tolerable thermal ranges. Although substantial variation exists within orders, Lepidoptera commonly exhibit behavioural and phenotypic buffering mechanisms-including microhabitat use, wing pigmentation, and phenological shifts-that reduce exposure to extreme temperatures (Rodrigues & Beldade, 2020). Similarly, many Hymenoptera, particularly ants, benefit from nesting strategies that exploit the thermal insulation of soil and shaded substrates. Ant colonies adjust nest depth and placement under rocks, logs, or cooler ground patches to regulate thermal exposure, thereby reducing vulnerability to elevated ambient temperatures (Parr & Bishop, 2022; Pincebourde & Suppo, 2016). The published thermal tolerance data used in our comparisons are based on relatively few taxa, with particularly limited information available for insects associated with oil palm systems. Consequently, these comparisons should be interpreted cautiously. Future studies should prioritise measuring the thermal tolerances of key oil palm species, including *Elaeidobius kamerunicus*, to improve predictions of climate vulnerability.

Temporal activity patterns further shape thermal risk. Insects operating during peak daytime temperatures may face greater exposure to thermal extremes, as peak fly activity in previous studies has been recorded to overlap with daily temperature maxima (López-Uribe et al., 2024), which are higher in the afternoon across both absolute ambient temperature and surface spatial characteristics in this study. In contrast, studies have found that bees tend to be more active during morning periods, which are generally cooler (López-Uribe et al., 2024). These diurnal differences could interact with microclimatic buffering to differentially influence thermal stress across taxa, underscoring the importance of structurally complex restored margins in moderating future climate impacts on insect communities by providing access to suitable thermal habitats throughout the day. In addition to exploiting cooler microhabitats, insects may also mitigate thermal stress through shifts in their diurnal activity patterns, a behavioural response documented under warming conditions (Franzén et al., 2022). Future research should therefore investigate how vegetation structural complexity and restoration management interact to influence both thermal refugia and behavioural thermoregulation in tropical agricultural landscapes, improving our understanding of species resilience to climate change.

Although Diptera were identified as being at elevated thermal risk across all restoration treatments and the control, these projections were derived from absolute ambient air temperatures and therefore do not capture fine-scale spatial variation in thermal conditions. Our surface temperature analyses indicate substantially greater thermal heterogeneity at finer spatial scales, where mosaics of hot and cool patches create opportunities for behavioural thermoregulation. Such microclimatic complexity can be ecologically important for mobile taxa, allowing individuals to exploit short-distance thermal refuges even when broader ambient conditions approach physiological limits (Scheffers et al., 2017). Consequently, models based solely on air temperature may overestimate thermal exposure for some taxa by overlooking the buffering capacity provided by structurally complex restored margins. By increasing the availability and accessibility of cool microhabitats throughout the day, especially under mature palm plantings, restoration may mitigate thermal stress and sustain insect activity, persistence, and functional roles under future climate warming.

### 4.5 Limitations, climate variability, and future directions

Interannual differences observed across the two field seasons, sampled in the dry and wet tropical climate, likely reflect broader ENSO-driven climate variability, which strongly influences rainfall, canopy moisture, and evapotranspiration dynamics across Southeast Asia (Cai et al., 2021). In particular, the first survey season coincided with the 2023–2024 El Niño, which brought warmer and generally drier-than-average conditions across much of Indonesia, whereas conditions returned closer to the long-term regional average during 2024 as ENSO weakened. Although these patterns are consistent with regional climate variability, our findings are necessarily constrained by the relatively small spatial extent of the study (two rivers) and two survey seasons. Given that tropical microclimates can vary substantially across landscapes and between years, longer-term monitoring across a greater number of sites is needed to more clearly separate restoration treatment effects from background climatic variation. Our projections further assume stationary microclimate offsets; however, buffering capacity may increase as canopies mature or decline if extreme heat or prolonged drought pushes vegetation toward physiological limits. Evaluating the persistence of microclimate benefits under compounded stressors, such as multi-year droughts, and explicitly quantifying cross-boundary thermal spillovers into adjacent crop areas represent key priorities for future research. Integrating structural restoration targets with long-term microclimate and biotic monitoring will be essential for improving management decisions and enhancing ecosystem resilience.

### 4.6 Management implications for climate-smart restoration and agricultural management

During the early restoration stage, we found a strong relationship between restoration treatment, structural development of riparian forests, and subsequent microclimatic conditions. Treatments that accelerated canopy closure and vertical layering, particularly enrichment planting combined with retained mature palms, delivered the most significant early microclimatic benefits, followed by a planting-only approach, which outperformed remnant mature palms and replanted young palms (control). These results highlight structural complexity as a key, tractable mechanism through which restoration can rapidly modify local thermal environments in agricultural landscapes.

Beyond short-term microclimatic moderation, these structural effects have important implications for the long-term persistence of biodiversity in human-modified tropical systems under climate change. In active oil palm plantations, many taxa are likely already operating in non-optimum conditions during peak heat periods under current climatic conditions, such that incremental warming or more frequent and severe heat extremes could rapidly push taxa to their thermal safety maximum. In this context, the capacity of restored riparian forests or broader management to increase canopy cover and reduce mean temperatures, dampen extremes, and lower thermal variability may be critical for preventing local population declines or extinction, particularly for thermally sensitive invertebrate communities.

Importantly, restoration not only reduced overall temperatures but also increased fine-scale thermal heterogeneity. Sensitivity varied among taxa; for groups with narrow thermal safety margins (e.g. some Diptera; López-Uribe et al., 2024; Yang et al., 1994), the cooler patches evident in actively restored plots may function as behavioural refuges during heat extremes. Such fine-scale heterogeneity is ecologically consequential, as mobile organisms can behaviourally thermoregulate by shifting among adjacent microhabitats to avoid thermal stress (Kearney et al., 2009; Pincebourde & Suppo, 2016). The persistence of these refugia may therefore play a key role in buffering biodiversity against increasing climatic variability in plantation landscapes.

Furthermore, thermal buffering is not uniform across forest strata. Above-ground environments experience greater exposure to solar radiation and more pronounced temperature fluctuations (e.g., canopy vs. ground), suggesting that taxa and ecological functions associated with these strata, such as pollination, biological control, and trophic regulation, may be particularly vulnerable in simplified agricultural systems (López-Uribe et al., 2024). This can have knock-on consequences when key taxa occupy specific strata within a forest structure, e.g., parasitoids in the canopy and their hosts in the understory, thereby knocking out key groups within a wider food web (Wenda et al., 2023). By accelerating canopy closure and promoting vertical structural complexity, active restoration could reduce thermal stress in these exposed strata, supporting not only species persistence but also the maintenance of ecosystem functions that contribute to agricultural productivity and resilience at plantation edges (Deutsch et al., 2018; Luke et al., 2019).

Together, these findings indicate that restoration and sustainability standards, such as those under the RSPO (RSPO, https://rspo.org/) and ISPO (ISPO, http://www.ispo-org.or.id), could benefit from moving beyond fixed riparian buffer widths to incorporate functional, structure-based targets. Metrics such as canopy cover thresholds, planting densities, and native species mixtures that are demonstrably linked to microclimate outcomes offer a climate-smart framework that could be achieved within the first decade of restoration efforts, while delivering co-benefits for biodiversity conservation and agricultural systems. More broadly, these findings highlight the importance of integrating restoration with plantation management practices. Reducing the extent of simultaneous canopy loss through approaches such as staggered replanting, combined with the strategic establishment of pioneer and later-successional tree species, could help maintain landscape-scale microclimatic buffering and habitat connectivity during plantation renewal. Such an approach would support the progressive development of forest structure while reducing thermal stress on biodiversity and enhancing the long-term resilience of both restored riparian margins and surrounding agricultural systems.

## 6. Acknowledgements

This work was funded by the John Stanley Gardiner Studentship Fund, Department of Zoology, University of Cambridge. We thank TEMPCON for supporting this research by providing discounted HOBO temperature loggers (UA-001-08 and U23-001). We thank Dr Gabriela Montejo-Kovacevich for her guidance on deploying the HOBO temperature loggers, Dr Michael D. Pashkevich for his assistance with terrestrial laser scanner fieldwork, and Dr Martin Ehbrecht for his expertise in processing the terrestrial laser data in FARO and RStudio. We thank PT Ivo Mas Tunggal, Golden Agri-Resources, and the SMART Research Institute (SMARTRI) for permission to conduct this research, access to the study sites, and logistical support throughout the RERTA project. We also thank the staff and field assistants at SMARTRI, Riau, for their invaluable assistance with fieldwork and data collection.

