## Supplementary Figure S1 for "From margins to mainstream: restoration of riparian margins to support oil palm sustainability in modern and future climates"

### Supplementary material

**Glossary T1.**

| **Term** | **Abbreviation** | **Definition** |
| --- | --- | --- |
| Terrestrial Lidar Scanner | TLS | Ground-based LiDAR scanner to measure vegetation structure. |
| Canopy Height | CH | Mean canopy height derived from TLS. |
| Canopy Openness | CO | Proportion of open sky seen from below. |
| Stand Structural Complexity Index | SSCI | Composite index of 3D forest structural complexity derived from TLS. |
| Effective Number of Layers | ENL | Vertical stratification measure representing canopy layering. |
| Mean Fractal Dimension | MeanFrac | Fractal descriptor related to canopy/structure complexity. |
| Vapour Pressure Deficit | VPD | Difference between saturation and actual vapour pressure (atmospheric drying power). |
| Diurnal Temperature Range | DTR | Daily maximum minus daily minimum temperature = DTR. |
| Land surface temperature | LST | Pixel temperatures of the forest floor. |
| Aggregation Index | AI | Proportion of shared edges among patches relative to the maximum possible; higher values indicate stronger clustering. |
| Hot patch / Cold patch area | Area | Mean area per hot/cold patch identified from thermal imagery. |
| Hot/Cold patch density | Density | Number of hot/cold patches per unit area. |
| Land Surface Temperature | LST | Temperature of the land surface estimated from thermal imagery. |
| Shape Index | SI | Patch shape metric; lower values indicate simpler (more compact) shapes. |
| Thermal Shannon Diversity Index | SHDI | Shannon diversity index applied to thermal classes; higher = more heterogeneous temperatures. |
| Neighbourhood contiguity | Queens contiguity | Spatial weights where neighbours share an edge or a corner (8-neighbour). |
| General Circulation Models | GCMs | Global climate models provide projections of future climate scenarios. |
| Shared Socioeconomic Pathways | SSPs | Greenhouse gas scenarios for future projections CHELSA. |
| Critical thermal maximum | CT_max | Upper thermal tolerance limit beyond which performance/viability declines sharply. |
| Proximity to CTmax | Proximity CTmax | Difference between CT_max and T_buffered,max._ |
| Buffered difference | TBuffer | Difference between buffered and ambient future temperatures. |
| Buffered future temperature | T_buffered,future_ | Projected ambient future temperature plus observed offset. |
| Temperature offset | Offset_observed_ | The difference between the margin (inside) and the outside (open) temperatures. |
| Thermal safety margin | TSM | Distance between environmental/body temperature and thermal limit. |

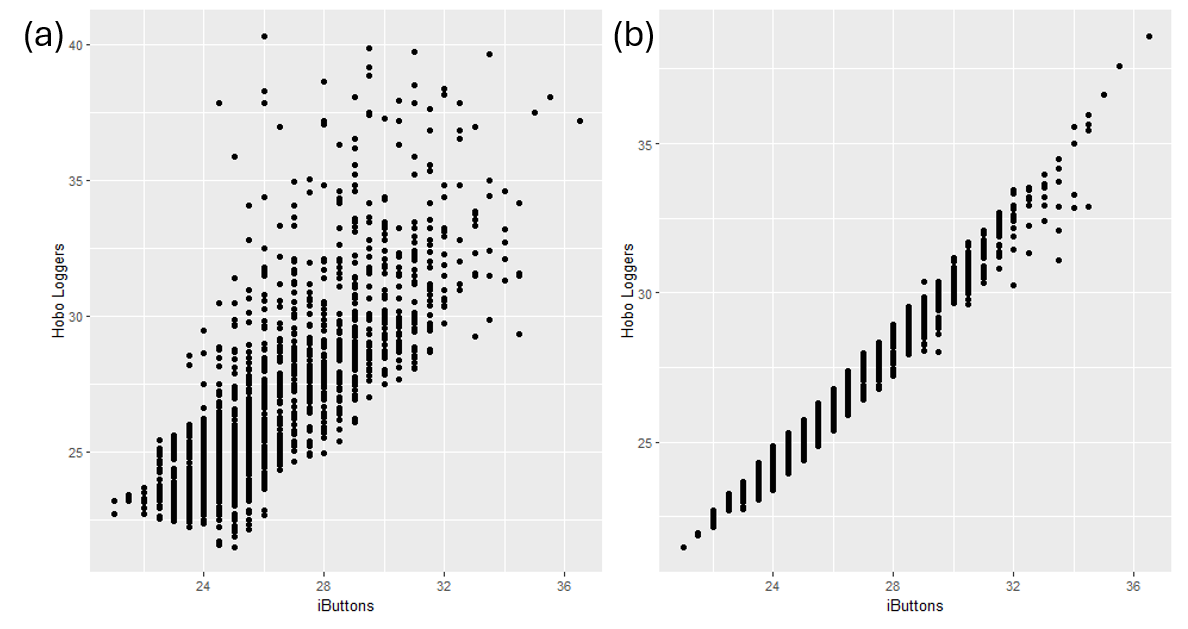
**Supplementary S1**:

**Figure S1:** Scatter plots showing the linear regression between temperatures recorded by two logger models: iButton (x-axis) and HOBO (y-axis). Both devices were co-located approximately 1.5 meters above ground and shielded from direct sunlight. Panel (a) shows data from the Dry season (2023); panel (b) shows data from the Wet season (2024). These plots were used to quantify systematic differences between the two device types, enabling correction of iButton temperature data for comparison across varying height measurements in subsequent analyses.

**Supplementary S2:**

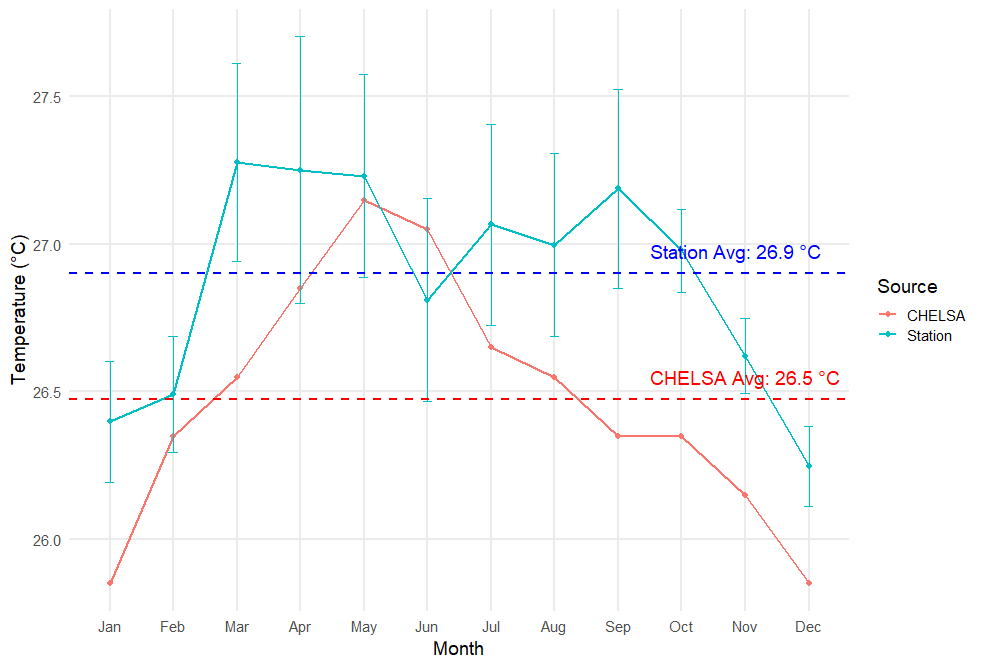

**Figure S2:** Comparison of monthly mean air temperatures derived from field-based weather station observations at the RERTA study sites (blue) and the CHELSA climatology (red), a high-resolution (~1 km) downscaled global climate dataset developed by the Swiss Federal Institute for Forest, Snow and Landscape Research (WSL). Monthly station temperatures represent the mean of the monthly averages calculated from observations collected between 2019 and 2024, whereas each CHELSA value represents a single long-term climatological monthly mean (1981–2000) for the corresponding location. Dashed horizontal lines indicate the overall annual mean temperature for each dataset (Station: 26.9 °C; CHELSA: 26.5 °C). Error bars are shown only for the weather station data and represent ±1 standard error (SE) among the yearly monthly means (2019–2024). Equivalent error bars are not shown for CHELSA because the extracted dataset provides one climatological monthly mean per month rather than annual replicate values. CHELSA consistently underestimated local temperatures relative to the field observations. This bias was quantified and subsequently used to bias-correct future CHELSA climate projections used in later analyses.

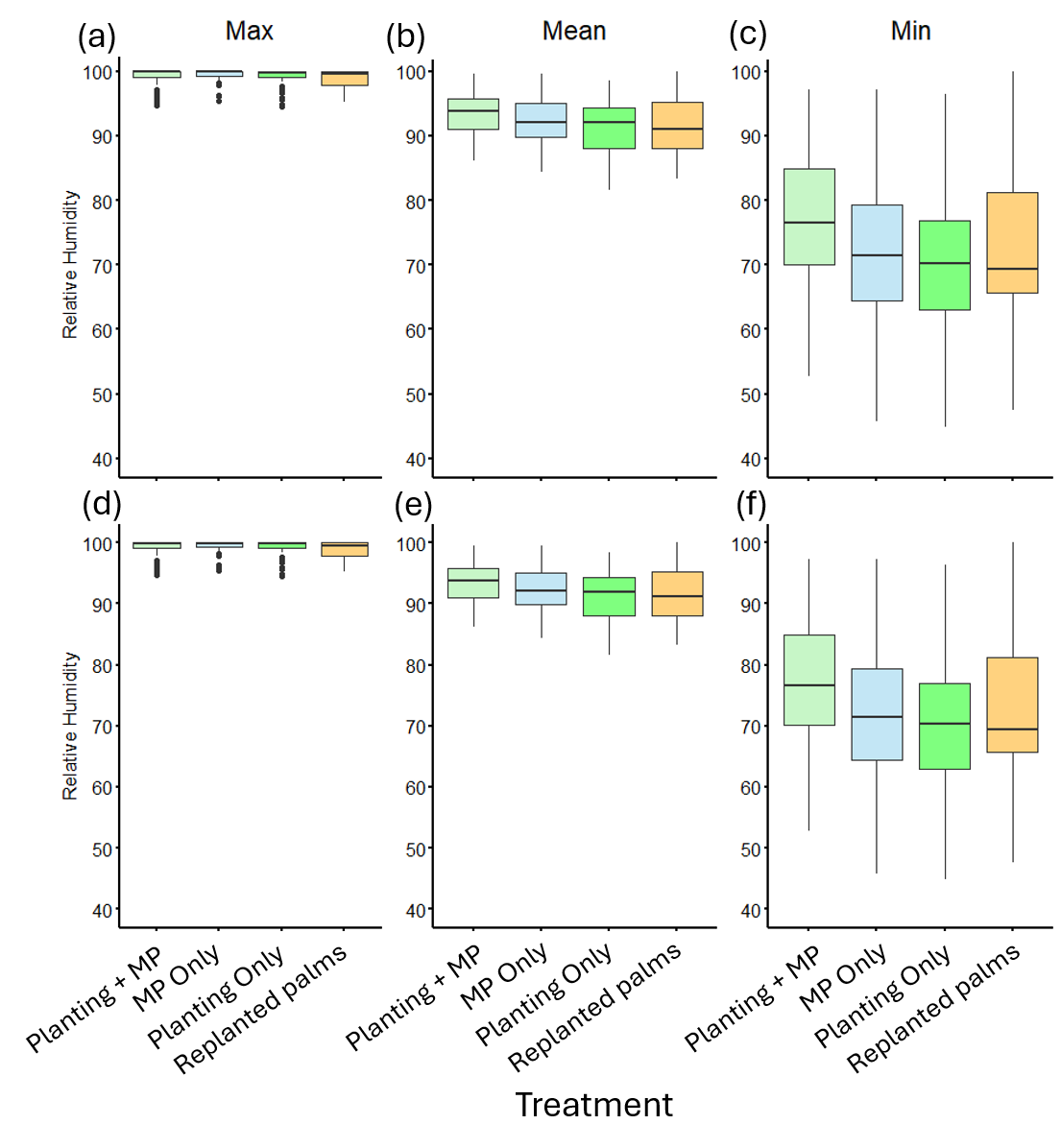
**Supplementary S3:**

**Figure S3:** Daily relative humidity recordings across forest heights (air, leaf litter and belowground) and restoration treatments in Dry (top row) and Wet season (bottom row). We first obtained individual datalogger daily maximum, mean, and minimum relative humidity, and then averaged these to obtain the daily mean values per treatment. Whiskers extend to the minimum and maximum values within 1.5 times the IQR. Points beyond this range are plotted as individual outliers.

| **Table S1:** The dataset is merged across multiple published thermal optimum (Topt), thermal safety margin (TSM), and critical thermal max (Ctmax) of tropical insects from published papers | | | | | | | | | |
| --- | --- | --- | --- | --- | --- | --- | --- | --- | --- |
| *Species* | Topt | Reference Topt | CTmax | TSM | Lat max | Long max | Reference max | Order | Family |
| *Drosophila birchii* |  |  | 35.5 |  | -17 | 146 | Hercus et al., 2000 | Diptera | Drosophilidae |
| *Drosophila serrata* |  |  | 36.7 |  | -17 | 146 | Hercus et al., 2000 | Diptera | Drosophilidae |
| *Aenictus sp.1* |  |  | 43.75 |  | 4.05 | 114.81 | Diamond et al., 2012 | Hymenoptera | Formicidae |
| *Anoplolepis gracilipes* |  |  | 45.25 |  | 4.05 | 114.81 | Diamond et al., 2012 | Hymenoptera | Formicidae |
| *Camponotus sp.1* | 34.1 | Bujan et al., 2016 | 46.25 | 12.15 | 4.05 | 114.81 | Diamond et al., 2012 | Hymenoptera | Formicidae |
| *Camponotus sp.2* | 34.1 | Bujan et al., 2016 | 45.5 | 11.4 | 4.05 | 114.81 | Diamond et al., 2012 | Hymenoptera | Formicidae |
| *Camponotus sp.3* | 34.1 | Bujan et al., 2016 | 45 | 10.9 | 4.05 | 114.81 | Diamond et al., 2012 | Hymenoptera | Formicidae |
| *Cardiocondyla sp.1* |  |  | 44.25 |  | 4.05 | 114.81 | Diamond et al., 2012 | Hymenoptera | Formicidae |
| *Cataulacus horridus* |  |  | 43.25 |  | 4.05 | 114.81 | Diamond et al., 2012 | Hymenoptera | Formicidae |
| *Crematogaster inflata* |  |  | 47.75 |  | 4.05 | 114.81 | Diamond et al., 2012 | Hymenoptera | Formicidae |
| *Crematogaster Physocremasp* |  |  | 40.75 |  | 4.05 | 114.81 | Diamond et al., 2012 | Hymenoptera | Formicidae |
| *Crematogaster sp.1* |  |  | 47.75 |  | 4.05 | 114.81 | Diamond et al., 2012 | Hymenoptera | Formicidae |
| *Crematogaster sp.2* |  |  | 46.75 |  | 4.05 | 114.81 | Diamond et al., 2012 | Hymenoptera | Formicidae |
| *Dolichoderus beccarii* |  |  | 45.63 |  | 4.05 | 114.81 | Diamond et al., 2012 | Hymenoptera | Formicidae |
| *Dolichoderus cuspidatus* |  |  | 44.5 |  | 4.05 | 114.81 | Diamond et al., 2012 | Hymenoptera | Formicidae |
| *GM-10-261* |  |  | 45.75 |  | 4.05 | 114.81 | Diamond et al., 2012 | Hymenoptera | Formicidae |
| *Gnamptogenys sp.1* |  |  | 43.75 |  | 4.05 | 114.81 | Diamond et al., 2012 | Hymenoptera | Formicidae |
| *Leptogenys sp.1* |  |  | 41.5 |  | 4.05 | 114.81 | Diamond et al., 2012 | Hymenoptera | Formicidae |
| *Monomorium floricola* |  |  | 47.5 |  | 4.05 | 114.81 | Diamond et al., 2012 | Hymenoptera | Formicidae |
| *Monomorium pharaonis* |  |  | 47.25 |  | 4.05 | 114.81 | Diamond et al., 2012 | Hymenoptera | Formicidae |
| *Myrmicaria sp.1* |  |  | 46 |  | 4.05 | 114.81 | Diamond et al., 2012 | Hymenoptera | Formicidae |
| *Pachycondyla tridentata* |  |  | 42 |  | 4.05 | 114.81 | Diamond et al., 2012 | Hymenoptera | Formicidae |
| *Pheidologeton sp.1* |  |  | 44.38 |  | 4.05 | 114.81 | Diamond et al., 2012 | Hymenoptera | Formicidae |
| *Polyrhachis bihamata* |  |  | 45 |  | 4.05 | 114.81 | Diamond et al., 2012 | Hymenoptera | Formicidae |
| *Polyrhachis furcata* |  |  | 44 |  | 4.05 | 114.81 | Diamond et al., 2012 | Hymenoptera | Formicidae |
| *Polyrhachis sp.1* |  |  | 43 |  | 4.05 | 114.81 | Diamond et al., 2012 | Hymenoptera | Formicidae |
| *Tapinoma sp.1* |  |  | 45.75 |  | 4.05 | 114.81 | Diamond et al., 2012 | Hymenoptera | Formicidae |
| *Technomyrmex sp.1* |  |  | 44.25 |  | 4.05 | 114.81 | Diamond et al., 2012 | Hymenoptera | Formicidae |
| *Tetramorium sp.1* |  |  | 44.7 |  | 4.05 | 114.81 | Diamond et al., 2012 | Hymenoptera | Formicidae |
| *Tetramorium Triglyphotrixsp* |  |  | 44.25 |  | 4.05 | 114.81 | Diamond et al., 2012 | Hymenoptera | Formicidae |
| *Messor denticornis* |  |  | 47.5 |  | -23.08 | 15.37 | Marsh, 1985 | Hymenoptera | Formicidae |
| *Monomorium alamarum* |  |  | 49 |  | -23.08 | 15.37 | Marsh, 1985 | Hymenoptera | Formicidae |
| *Monomorium vatranum* |  |  | 49 |  | -23.08 | 15.37 | Marsh, 1985 | Hymenoptera | Formicidae |
| *Monomorium viator* |  |  | 46.7 |  | -23.08 | 15.37 | Marsh, 1985 | Hymenoptera | Formicidae |
| *Ocymyrmex barbiger* |  |  | 51.4 |  | -23.08 | 15.37 | Marsh, 1985 | Hymenoptera | Formicidae |
| *Pheidole tenuinodis* |  |  | 42.9 |  | -23.08 | 15.37 | Marsh, 1985 | Hymenoptera | Formicidae |
| *Tetramorium rufescens* |  |  | 46.9 |  | -23.08 | 15.37 | Marsh, 1985 | Hymenoptera | Formicidae |
| *Tetramorium sericeiventre* |  |  | 47.9 |  | -23.08 | 15.37 | Marsh, 1985 | Hymenoptera | Formicidae |
| *Camponotus detritus* |  |  | 53.8 |  | -23.08 | 15.37 | Marsh, 1985 | Hymenoptera | Formicidae |
| *Pogonomyrmex desertorum* |  |  | 53.3 |  | -23.08 | 15.37 | Marsh, 1985 | Hymenoptera | Formicidae |
| *Epigomphus quadracies* |  |  | 41.1 |  | 9.16 | -79.83 | May, 1976 | Odonata | Gomphidae |
| *Erythemis credula* |  |  | 45.6 |  | 9.16 | -79.83 | May, 1976 | Odonata | Libellulidae |
| *Erythemis plebeja* |  |  | 46.8 |  | 9.16 | -79.83 | May, 1976 | Odonata | Libellulidae |
| *Miathyria marcella* |  |  | 47 |  | 9.16 | -79.83 | May, 1976 | Odonata | Libellulidae |
| *Micrathyria aequalis* |  |  | 45.4 |  | 9.16 | -79.83 | May, 1976 | Odonata | Libellulidae |
| *Micrathyria eximia* |  |  | 44.7 |  | 9.16 | -79.83 | May, 1976 | Odonata | Libellulidae |
| *Micrathyria ocellata* |  |  | 44.3 |  | 9.16 | -79.83 | May, 1976 | Odonata | Libellulidae |
| *Tauriphila argo* |  |  | 48.5 |  | 9.16 | -79.83 | May, 1976 | Odonata | Libellulidae |
| *Tramea cophysa* |  |  | 47.6 |  | 9.16 | -79.83 | May, 1976 | Odonata | Libellulidae |
| *Tramea walkeri* |  |  | 46.3 |  | 9.16 | -79.83 | May, 1976 | Odonata | Libellulidae |
| *Glossina pallidipes* |  |  | 44.3 |  | -13 | 31.5 | Terblanche et_al., 2008 | Diptera | Glossinidae |
| *Adelpha cytherea* |  |  | 48.07 |  | 9.116 | -79.697 | Ashe-Jepson et al., 2023 | Lepidoptera | Nymphalidae |
| *Anartia fatima* |  |  | 49.43 |  | 9.116 | -79.697 | Ashe-Jepson et al., 2023 | Lepidoptera | Nymphalidae |
| *Anartia jatrophae* |  |  | 53.3 |  | 9.116 | -79.697 | Ashe-Jepson et al., 2023 | Lepidoptera | Nymphalidae |
| *Arawacus togarna* |  |  | 49.76 |  | 9.116 | -79.697 | Ashe‐Jepson et al., 2023 | Lepidoptera | Lycaenidae |
| *Calephelis sp* |  |  | 50.7 |  | 9.116 | -79.697 | Ashe‐Jepson et al., 2023 | Lepidoptera | Riodinidae |
| *Calycopis isobeon* |  |  | 47.8 |  | 9.116 | -79.697 | Ashe‐Jepson et al., 2023 | Lepidoptera | Lycaenidae |
| *Cissa pompilia* |  |  | 49.94 |  | 9.116 | -79.697 | Ashe‐Jepson et al., 2023 | Lepidoptera | Nymphalidae |
| *Cissia pseudoconfusa* |  |  | 49.91 |  | 9.116 | -79.697 | Ashe‐Jepson et al., 2023 | Lepidoptera | Nymphalidae |
| *Dione juno* |  |  | 45.53 |  | 9.116 | -79.697 | Ashe‐Jepson et al., 2023 | Lepidoptera | Nymphalidae |
| *Eurema daira* |  |  | 52.31 |  | 9.116 | -79.697 | Ashe‐Jepson et al., 2023 | Lepidoptera | Pieridae |
| *Hemiargus hanno* |  |  | 51.27 |  | 9.116 | -79.697 | Ashe‐Jepson et al., 2023 | Lepidoptera | Lycaenidae |
| *Hermeuptychia hermes* | 26 |  | 51.01 |  | 9.116 | -79.697 | Ashe‐Jepson et al., 2023 | Lepidoptera | Nymphalidae |
| *Hylephila phyleus* | 35 | Tashiro & Mitchell, 1985 | 51.16 | 16.16 | 9.116 | -79.697 | Ashe‐Jepson et al., 2023 | Lepidoptera | Hesperiidae |
| *Itaballia demophile* |  |  | 45.97 |  | 9.116 | -79.697 | Ashe‐Jepson et al., 2023 | Lepidoptera | Pieridae |
| *Juditha caucana* |  |  | 46.23 |  | 9.116 | -79.697 | Ashe‐Jepson et al., 2023 | Lepidoptera | Riodinidae |
| *Junonia_sp1* |  |  | 55.24 |  | 9.116 | -79.697 | Ashe‐Jepson et al., 2023 | Lepidoptera | Nymphalidae |
| *Nica flavilla* |  |  | 47.67 |  | 9.116 | -79.697 | Ashe‐Jepson et al., 2023 | Lepidoptera | Nymphalidae |
| *Phoebis sennae* |  |  | 49.59 |  | 9.116 | -79.697 | Ashe‐Jepson et al., 2023 | Lepidoptera | Pieridae |
| *Pompeius pompeius* |  |  | 51.71 |  | 9.116 | -79.697 | Ashe‐Jepson et al., 2023 | Lepidoptera | Hesperiidae |
| *Pyrgus orcas* |  |  | 51.92 |  | 9.116 | -79.697 | Ashe‐Jepson et al., 2023 | Lepidoptera | Hesperiidae |
| *Staphylus vulgata* |  |  | 47.74 |  | 9.116 | -79.697 | Ashe‐Jepson et al., 2023 | Lepidoptera | Hesperiidae |
| *Urbanus procne* |  |  | 51.72 |  | 9.116 | -79.697 | Ashe‐Jepson et al., 2023 | Lepidoptera | Hesperiidae |
| *Urbanus simplicius* |  |  | 51.9 |  | 9.116 | -79.697 | Ashe‐Jepson et al., 2023 | Lepidoptera | Hesperiidae |
| *Vehilius stictomenes* |  |  | 51.83 |  | 9.116 | -79.697 | Ashe‐Jepson et al., 2023 | Lepidoptera | Hesperiidae |
| *Bactrocera cucurbitae* | 24.7 | Yang et al., 1994b | 29.18 | 4.48 | 16.3 | 111.4 | Yang et al., 1994b | Diptera |  |
| *Bactrocera latifrons* | 28.4 | Vargas et al., 1997 | 32.88 | 4.48 | 21.5 | -158 | Vargas et al., 1997 | Diptera |  |
| *Ceratitis capitata* | 27.98 | Vargas et al., 1997 | 31.78 | 3.8 | 20.88 | -156.52 | Vargas et al., 1997 | Diptera |  |
| *Bactrocera dorsalis* | 25 | Michel A. et al., 2021 | 36.2 | 11.2 | 23.12 | 113.25 | Yan et al., 1994 | Diptera |  |
| *Clavigralla shadabi* | 28.8 | Dreyer & Baumgärtner, 1996 | 33.28 | 4.48 | 6.5 | 2.4 | Dreyer & Baumgärtner, 1996 | Hemiptera |  |
| *Clavigralla tomentosicollis* | 32.6 | Dreyer & Baumgärtner, 1996 | 37.08 | 4.48 | 6.5 | 2.4 | Dreyer & Baumgärtner, 1996 | Hemiptera |  |
| *Saccharicoccus sacchari* | 30.68 | Rae & De’Ath, 1991 | 34.78 | 4.1 | -21.09 | 149.09 | Rae & De’Ath, 1991 | Hemiptera |  |
| *Clavigralla shadabi* | 28.8 | Dreyer & Baumgärtner, 1996 | 33.0677 | 4.2677 | 6.45 | 2.35 | Dreyer & Baumgärtner, 1996 | Hemiptera |  |
| *Cotesia sesamiae* | 30.05 | Mbapila and Overholt, 2001 | 36.65 | 6.6 | -3 | 40 | Mbapila & Overholt, 2001 | Hymenoptera |  |
| *Uscana lariophaga* | 36.3 | van Huis et al., 1994 | 41.04 | 4.74 | 13.48 | 2.17 | van Huis et al., 1994 | Hymenoptera |  |

**Table S1:** Max, Mean, and Min temperatures across the restoration treatments (Planting + MP: enrichment planting with mature oil palm; MP only: mature oil palm only; Planting only: mature oil palm removed with enrichment planting; Palms replanted: control— oil palms planted to river edge) in Dry (2023) and Wet (2024). We estimated the daily maximum, mean, and minimum temperatures (◦C) per data logger per day, and averaged across the recorded time (~14 days). Values shown are mean ± s.e.m.

| Season | Treatment | T_max_ Air | T_max_ Soil | T_max_ Leaf Litter | T_mean_ Air | T_mean_ Soil | T_mean_ Leaf Litter | T_min_ Air | T_min_ Soil | T_min_ Leaf Litter |
| --- | --- | --- | --- | --- | --- | --- | --- | --- | --- | --- |
| Dry | Planting + MP | 31.2 ± 0.3 | 28.8 ± 0.29 | 28.1 ± 0.15 | 24.9 ± 0.1 | 26.5 ± 0.21 | 26.1 ± 0.17 | 24.9 ± 0.1 | 26.5 ± 0.21 | 26.1 ± 0.17 |
| Dry | MP only | 33.3 ± 0.33 | 28.1 ± 0.11 | 28 ± 0.12 | 24.6 ± 0.11 | 26.5 ± 0.16 | 25.2 ± 0.18 | 24.6 ± 0.11 | 26.5 ± 0.16 | 25.2 ± 0.18 |
| Dry | Planting only | 32 ± 0.26 | 28.1 ± 0.08 | 29.1 ± 0.15 | 24.6 ± 0.15 | 26.8 ± 0.12 | 26.5 ± 0.18 | 24.6 ± 0.15 | 26.8 ± 0.12 | 26.5 ± 0.18 |
| Dry | Replanted palms | 35.4 ± 0.44 | 29 ± 0.11 | 32 ± 0.28 | 24.2 ± 0.17 | 26.7 ± 0.4 | 25.6 ± 0.27 | 24.2 ± 0.17 | 26.7 ± 0.4 | 25.6 ± 0.27 |
| Wet | Planting + MP | 30.7 ± 0.29 | 27.9 ± 0.12 | 28.7 ± 0.13 | 25.1 ± 0.05 | 26.4 ± 0.11 | 26.3 ± 0.2 | 25.1 ± 0.05 | 26.7 ± 0.11 | 26.6 ± 0.2 |
| Wet | MP only | 33.2 ± 0.47 | 28.8 ± 0.21 | 31 ± 0.26 | 25 ± 0.08 | 26.4 ± 0.15 | 25.7 ± 0.27 | 25 ± 0.08 | 26.6 ± 0.15 | 25.9 ± 0.27 |
| Wet | Planting only | 32.2 ± 0.32 | 29.3 ± 0.2 | 30 ± 0.2 | 25.1 ± 0.06 | 26.6 ± 0.18 | 25.8 ± 0.2 | 25.1 ± 0.06 | 26.8 ± 0.18 | 26.1 ± 0.2 |
| Wet | Replanted palms | 32.7 ± 0.34 | 29 ± 0.19 | 29 ± 0.13 | 24.9 ± 0.07 | 26.8 ± 0.14 | 26.2 ± 0.1 | 24.9 ± 0.07 | 27 ± 0.14 | 26.4 ± 0.1 |

**Table S2:** Riparian margin temperature offsets across the restoration treatments (Planting + MP: enrichment planting with mature oil palm; MP only: mature oil palm only; Planting only: mature oil palm removed with enrichment planting; Palms replanted: control— oil palms planted to river edge) and field seasons (Dry 2023; Wet 2024).

| Year | Site | Treatment | T_max_ Offset | T_mean_ Offset | T_min_ Offset |
| --- | --- | --- | --- | --- | --- |
| Dry | River 1 | Planting + MP | -2.31 ± 0.304 | -0.59 ± 0.098 | 0.51 ± 0.095 |
| Dry | River 1 | MP only | 3.84 ± 0.456 | 0.19 ± 0.041 | 0.32 ± 0.081 |
| Dry | River 1 | Planting only | 0.56 ± 0.314 | -0.29 ± 0.063 | 0.32 ± 0.091 |
| Dry | River 1 | Replanted palms | 4.79 ± 0.525 | 0.31 ± 0.048 | 0.01 ± 0.094 |
| Dry | River 2 | Planting + MP | 3.67 ± 0.547 | -0.02 ± 0.025 | 0.14 ± 0.086 |
| Dry | River 2 | MP only | 5.1 ± 0.726 | 0.04 ± 0.038 | -0.04 ± 0.076 |
| Dry | River 2 | Planting only | 2.41 ± 0.298 | -0.08 ± 0.03 | -0.22 ± 0.068 |
| Dry | River 2 | Replanted palms | 7.82 ± 0.833 | 1.17 ± 0.173 | -0.43 ± 0.086 |
| Wet | River 1 | Planting + MP | -2.43 ± 0.295 | -0.72 ± 0.089 | 0.22 ± 0.121 |
| Wet | River 1 | MP only | 6.86 ± 0.799 | 0.04 ± 0.086 | 0.1 ± 0.126 |
| Wet | River 1 | Planting only | 0.2 ± 0.256 | -0.25 ± 0.072 | 0.22 ± 0.128 |
| Wet | River 1 | Replanted palms | 2.56 ± 0.374 | -0.01 ± 0.086 | 0.01 ± 0.128 |
| Wet | River 2 | Planting + MP | 1.38 ± 0.41 | -0.31 ± 0.055 | -0.02 ± 0.095 |
| Wet | River 2 | MP only | 3.59 ± 0.669 | -0.13 ± 0.064 | -0.07 ± 0.093 |
| Wet | River 2 | Planting only | 2.62 ± 0.381 | -0.05 ± 0.067 | -0.36 ± 0.092 |
| Wet | River 2 | Replanted palms | 3.11 ± 0.408 | 0.09 ± 0.084 | -0.58 ± 0.1 |

**Supplementary Table S3:** Post Hoc Comparisons for Vegetative Structural Complexity metrics measured using the Terrestrial Lidar Scanner (Stand Structural Complexity Index; Enhanced Layering; MeanFrac; Canopy Openness; Canopy Height) between restoration treatments (Planting + MP: enrichment planting with mature oil palm; MP only: mature oil palm only; Planting only: mature oil palm removed with enrichment planting; Palms replanted: control— oil palms planted to river edge).

| Post Hoc Comparisons for Vegetative Structural Complexity Between Restoration Treatments | | | | | | | |
| --- | --- | --- | --- | --- | --- | --- | --- |
| Metric | Comparison | Difference | SE | df | Test Statistic | p-value | Test |
| Stand Structural Complexity Index (SSCI) | Planting + MP – MP only | -2.23 | - |  | - | 0.0028 | Tukey HSD |
|  | Planting + MP – Planting only | 0.80 | - |  | - | 0.5042 | Tukey HSD |
|  | Planting + MP – Replanted palms | -2.07 | - |  | - | 0.0058 | Tukey HSD |
|  | MP only – Planting only | 3.04 | - |  | - | <0.0001 | Tukey HSD |
|  | MP only – Replanted palms | 0.16 | - |  | - | 0.9919 | Tukey HSD |
|  | Planting only – Replanted palms | -2.88 | - |  | - | <0.0002 | Tukey HSD |
| Metric | Comparison |  |  |  | Test Statistic | p-value | Test |
| Enhanced Layering (ENL) | Planting + MP – MP only | - | - | - | 0.13 | 1 | Dunn |
|  | Planting + MP – Planting only | - | - | - | 0.19 | 1 | Dunn |
|  | Planting + MP – Replanted palms | - | - | - | 3.52 | 0.0013 | Dunn |
|  | MP only – Planting only | - | - | - | 0.05 | 1 | Dunn |
|  | MP only –Replanted palms | - | - | - | 3.38 | 0.0021 | Dunn |
|  | Planting only – Replanted palms | - | - | - | 3.33 | 0.0026 | Dunn |
| Metric | Comparison | Estimate | SE | df | t-ratio | p-value | Test |
| MeanFrac | Planting + MP – MP only | 0.2266 | 0.0655 | 27 | 3.461 | 0.0092 | EMMs |
|  | Planting + MP – Planting only | -0.0639 | 0.0655 | 27 | -0.975 | 0.7645 | EMMs |
|  | Planting + MP – Replanted palms | -0.1189 | 0.0655 | 27 | -1.816 | 0.2881 | EMMs |
|  | MP only – Planting only | -0.2905 | 0.0655 | 27 | -4.436 | 0.0008 | EMMs |
|  | MP only – Replanted palms | -0.3455 | 0.0655 | 27 | -5.277 | 0.0001 | EMMs |
|  | Planting only – Replanted palms | -0.0550 | 0.0655 | 27 | -0.840 | 0.8348 | EMMs |
| Metric | Comparison | Estimate | SE | df | z-ratio | p-value | Test type |
| Canopy Openness | Planting + MP – MP only | -3.884 | 1.32 | Inf | -2.949 | 0.0168 | EMMs |
|  | Planting + MP – Planting only | -0.313 | 1.32 | Inf | -0.237 | 0.9953 | EMMs |
|  | Planting + MP – Replanted palms | -4.841 | 1.32 | Inf | -3.675 | 0.0014 | EMMs |
|  | MP only – Planting only | 3.572 | 1.31 | Inf | 2.729 | 0.0322 | EMMs |
|  | MP only – Replanted palms | -0.957 | 1.3 | Inf | -0.735 | 0.8831 | EMMs |
|  | Planting only – Replanted palms | -4.528 | 1.31 | Inf | -3.460 | 0.0030 | EMMs |
| Metric | Comparison |  |  |  | Test Statistic | p-value | Test |
| Canopy Height | Planting + MP – MP only | - | - | - | 0.83 | 1 | Dunn |
|  | Planting + MP – Planting only | - | - | - | 0.78 | 1 | Dunn |
|  | Planting + MP – Replanted palms | - | - | - | -0.05 | 1 | Dunn |
|  | MP only – Planting only | - | - | - | 3.97 | 0.0002 | Dunn |
|  | MP only – Replanted palms | - | - | - | 3.14 | 0.0051 | Dunn |
|  | Planting only – Replanted palms | - | - | - | 3.19 | 0.0042 | Dunn |

**Supplementary Table S4:** Post Hoc comparisons for the temperature recorded at different heights (Macroclimate = air temperature; Leaf litter = ~ 5cm depth; below ground ~ 5cm depth) between treatments (Planting + MP: enrichment planting with mature oil palm; MP only: mature oil palm only; Planting only: mature oil palm removed with enrichment planting; Palms replanted: control— oil palms planted to river edge).

| Post Hoc comparisons for the temperature recorded at different heights between treatments | | | | | | | | |
| --- | --- | --- | --- | --- | --- | --- | --- | --- |
| Dry Season (2023: July) | | | | | | | | |
| Metric | Type | Comparison | Estimate | SE | df | Test ratio | p-value | Test |
| Macroclimate (~1.5 m aboveground) | Max | Planting + MP – MP only | -2.060 | 1.19 | 35.6 | -1.736 | 0.3208 | EMMs |
|  |  | Planting + MP – Planting only | -0.816 | 1.19 | 35.6 | -0.687 | 0.9012 | EMMs |
|  |  | Planting + MP – Replanted palms | -4.191 | 1.23 | 35.6 | -3.412 | 0.0084 | EMMs |
|  |  | MP only – Planting only | 1.244 | 1.19 | 35.6 | 1.048 | 0.7225 | EMMs |
|  |  | MP only – Replanted palms | -2.131 | 1.23 | 35.6 | -1.735 | 0.3211 | EMMs |
|  |  | Planting only – Replanted palms | -3.375 | 1.23 | 35.6 | -2.748 | 0.0442 | EMMs |
|  | Mean | Planting + MP – MP only | -0.419 | 0.254 | 35.6 | -1.650 | 0.3648 | EMMs |
|  |  | Planting + MP – Planting only | -0.123 | 0.254 | 35.6 | -0.485 | 0.9619 | EMMs |
|  |  | Planting + MP – Replanted palms | -0.986 | 0.263 | 35.6 | -3.748 | 0.0034 | EMMs |
|  |  | MP only – Planting only | 0.296 | 0.254 | 35.6 | 1.165 | 0.6525 | EMMs |
|  |  | MP only – Replanted palms | -0.567 | 0.263 | 35.6 | -2.154 | 0.1561 | EMMs |
|  |  | Planting only – Replanted palms | -0.863 | 0.263 | 35.6 | -3.279 | 0.0119 | EMMs |
|  | Minimum | Planting + MP – MP only | 0.1923 | 0.144 | 35.6 | 1.334 | 0.5482 | EMMs |
|  |  | Planting + MP – Planting only | 0.2233 | 0.144 | 35.6 | 1.549 | 0.4199 | EMMs |
|  |  | Planting + MP – Replanted palms | 0.4056 | 0.149 | 35.6 | 2.718 | 0.0474 | EMMs |
|  |  | MP only – Planting only | 0.0311 | 0.144 | 35.6 | 0.215 | 0.9964 | EMMs |
|  |  | MP only – Replanted palms | 0.2133 | 0.149 | 35.6 | 1.429 | 0.4901 | EMMs |
|  |  | Planting only – Replanted palms | 0.1822 | 0.149 | 35.6 | 1.221 | 0.6179 | EMMs |
| Metric | Type | Comparison | Estimate | SE | df | Test ratio | p-value | Test |
| Leaf litter (~5 cm) | Max | Planting + MP – MP only | 0.0052 | 0.0396 | 19.6 | 0.131 | 0.9992 | EMMs |
|  |  | Planting + MP – Planting only | -0.0361 | 0.0366 | 19.6 | -0.986 | 0.7590 | EMMs |
|  |  | Planting + MP – Replanted palms | -0.1264 | 0.0396 | 19.6 | -3.196 | 0.0220 | EMMs |
|  |  | MP only – Planting only | -0.0413 | 0.0396 | 19.6 | -1.044 | 0.7262 | EMMs |
|  |  | MP only – Replanted palms | -0.1316 | 0.0423 | 19.6 | -3.112 | 0.0263 | EMMs |
|  |  | Planting only – Replanted palms | -0.0903 | 0.0396 | 19.6 | -2.283 | 0.1364 | EMMs |
|  | Mean | Planting + MP – MP only | 0.448 | 0.460 | 19.6 | 0.974 | 0.7654 | EMMs |
|  |  | Planting + MP – Replanted palms | -0.602 | 0.462 | 19.6 | -1.412 | 0.5068 | EMMs |
|  |  | MP only – Planting only | -0.930 | 0.460 | 19.6 | -2.020 | 0.2147 | EMMs |
|  |  | MP only – Planting only | -1.050 | 0.460 | 19.6 | -2.282 | 0.1366 | EMMs |
|  |  | MP only – Replanted palms | -1.378 | 0.492 | 19.6 | -2.801 | 0.0503 | EMMs |
|  |  | Planting only – Replanted palms | -0.328 | 0.460 | 19.6 | -0.712 | 0.8911 | EMMs |
|  | Minimum | Planting + MP – MP only | 1.044 | 0.414 | 19.6 | 2.526 | 0.0867 | EMMs |
|  |  | Planting + MP – Planting only | -0.342 | 0.383 | 19.6 | -0.892 | 0.8088 | EMMs |
|  |  | Planting + MP – Replanted palms | 0.389 | 0.414 | 19.6 | 0.940 | 0.7837 | EMMs |
|  |  | MP only – Planting only | -1.386 | 0.414 | 19.6 | -3.352 | 0.0157 | EMMs |
|  |  | MP only – Replanted palms | -0.656 | 0.442 | 19.6 | -1.483 | 0.4660 | EMMs |
|  |  | Planting only – Replanted palms | 0.731 | 0.414 | 19.6 | 1.767 | 0.3182 | EMMs |
| Metric |  | Comparison | Estimate | SE | df | t-ratio | p-value | Test |
| Below ground (~5 cm) | Max | Planting + MP – MP only | 0.6542 | 0.758 | 34.6 | 0.864 | 0.8235 | EMMs |
|  |  | Planting + MP – Planting only | 0.6655 | 0.784 | 34.6 | 0.849 | 0.8308 | EMMs |
|  |  | Planting + MP – Replanted palms | -0.1631 | 0.784 | 34.6 | -0.208 | 0.9968 | EMMs |
|  |  | MP only – Planting only | 0.0113 | 0.784 | 34.6 | 0.014 | 1 | EMMs |
|  |  | MP only – Replanted palms | -0.8173 | 0.784 | 34.6 | -1.042 | 0.7261 | EMMs |
|  |  | Planting only – Replanted palms | -0.8286 | 0.810 | 34.6 | -1.023 | 0.7372 | EMMs |
|  | Mean | Planting + MP – MP only | -0.0279 | 0.246 | 34.6 | -0.113 | 0.9995 | EMMs |
|  |  | Planting + MP – Planting only | -0.2607 | 0.254 | 34.6 | -1.025 | 0.7362 | EMMs |
|  |  | Planting + MP – Replanted palms | -0.4010 | 0.254 | 34.6 | -1.576 | 0.4051 | EMMs |
|  |  | MP only – Planting only | -0.2328 | 0.254 | 34.6 | -0.915 | 0.7969 | EMMs |
|  |  | MP only – Replanted palms | -0.3731 | 0.254 | 34.6 | -1.467 | 0.4680 | EMMs |
|  |  | Planting only – Replanted palms | -0.1403 | 0.263 | 34.6 | -0.534 | 0.9502 | EMMs |
|  | Minimum | Planting + MP – MP only | -0.154 | 0.365 | 34.6 | -0.423 | 0.9742 | EMMs |
|  |  | Planting + MP – Planting only | -0.522 | 0.377 | 34.6 | -1.383 | 0.5181 | EMMs |
|  |  | Planting + MP – Replanted palms | -0.289 | 0.377 | 34.6 | -0.765 | 0.8696 | EMMs |
|  |  | MP only – Planting only | -0.368 | 0.377 | 34.6 | -0.975 | 0.7646 | EMMs |
|  |  | MP only – Replanted palms | 0.135 | 0.377 | 34.6 | -0.356 | 0.9842 | EMMs |
|  |  | Planting only – Replanted palms | 0.233 | 0.390 | 34.6 | 0.599 | 0.9318 | EMMs |
| Wet Season (2024: February) | | | | | | | | |
| Metric | Type | Comparison | Estimate | SE | df | t-ratio | p-value | Test |
| Macroclimate (~1.5 m aboveground) | Max | Planting + MP – MP only | -2.461 | 1.07 | 36.6 | -2.294 | 0.1180 | EMMs |
|  |  | Planting + MP – Planting only | -1.436 | 1.07 | 36.6 | -1.339 | 0.5448 | EMMs |
|  |  | Planting + MP – Replanted palms | -1.982 | 1.07 | 36.6 | -1.847 | 0.2683 | EMMs |
|  |  | MP only – Planting only | 1.025 | 1.07 | 36.6 | 0.955 | 0.7753 | EMMs |
|  |  | MP only – Replanted palms | 0.479 | 1.07 | 36.6 | 0.447 | 0.9698 | EMMs |
|  |  | Planting only – Replanted palms | -0.545 | 1.07 | 36.6 | -0.509 | 0.9565 | EMMs |
|  | Mean | Planting + MP – MP only | -0.419 | 0.254 | 35.6 | -1.650 | 0.3648 | EMMs |
|  |  | Planting + MP – Planting only | -0.123 | 0.254 | 35.6 | -0.485 | 0.9619 | EMMs |
|  |  | Planting + MP – Replanted palms | -0.986 | 0.263 | 35.6 | -3.748 | 0.0034 | EMMs |
|  |  | MP only – Planting only | 0.296 | 0.254 | 35.6 | 1.165 | 0.6525 | EMMs |
|  |  | MP only – Replanted palms | -0.567 | 0.263 | 35.6 | -2.154 | 0.1561 | EMMs |
|  |  | Planting only – Replanted palms | 0.863 | 0.263 | 35.6 | -3.279 | 0.0119 | EMMs |
|  | Minimum | Planting + MP – MP only | 0.1923 | 0.144 | 35.6 | 1.334 | 0.5482 | EMMs |
|  |  | Planting + MP – Planting only | 0.2233 | 0.144 | 35.6 | 1.549 | 0.4199 | EMMs |
|  |  | Planting + MP – Replanted palms | 0.4056 | 0.149 | 35.6 | 2.718 | 0.0474 | EMMs |
|  |  | MP only – Planting only | 0.0311 | 0.144 | 35.6 | 0.215 | 0.9964 | EMMs |
|  |  | MP only – Replanted palms | 0.2133 | 0.149 | 35.6 | 1.429 | 0.4901 | EMMs |
|  |  | Planting only – Replanted palms | 0.1822 | 0.149 | 35.6 | 1.221 | 0.6179 | EMMs |
| Metric | Type | Comparison | Estimate | SE | df | t-ratio | p-value | Test |
| Below ground (~5 cm) | Max | Planting + MP – MP only | -0.885 | 0.814 | 32.7 | -1.087 | 0.6999 | EMMs |
|  |  | Planting + MP – Planting only | -1.381 | 0.714 | 32.7 | -1.933 | 0.2345 | EMMs |
|  |  | Planting + MP – Replanted palms | -1.131 | 0.739 | 32.7 | -1.529 | 0.4323 | EMMs |
|  |  | MP only – Planting only | -0.495 | 0.814 | 32.7 | -0.608 | 0.9287 | EMMs |
|  |  | MP only – Replanted palms | -0.245 | 0.837 | 32.7 | -0.293 | 0.9910 | EMMs |
|  |  | Planting only – Replanted palms | 0.250 | 0.739 | 32.7 | 0.338 | 0.9864 | EMMs |
|  | Mean | Planting + MP – MP only | -0.0279 | 0.246 | 34.6 | -0.113 | 0.9995 | EMMs |
|  |  | Planting + MP – Planting only | -0.2607 | 0.254 | 34.6 | -1.025 | 0.7362 | EMMs |
|  |  | Planting + MP – Replanted palms | -0.4010 | 0.254 | 34.6 | -1.576 | 0.4051 | EMMs |
|  |  | MP only – Planting only | -0.2328 | 0.254 | 34.6 | -0.915 | 0.7969 | EMMs |
|  |  | MP only – Replanted palms | -0.3731 | 0.254 | 34.6 | -1.467 | 0.4680 | EMMs |
|  |  | Planting only – Replanted palms | -0.1403 | 0.263 | 34.6 | -0.534 | 0.9502 | EMMs |
|  | Minimum | Planting + MP – MP only | -0.133 | 0.360 | 34.6 | -0.370 | 0.9824 | EMMs |
|  |  | Planting + MP – Planting only | -0.511 | 0.373 | 34.6 | -1.370 | 0.5260 | EMMs |
|  |  | Planting + MP – Replanted palms | -0.282 | 0.373 | 34.6 | -0.757 | 0.8731 | EMMs |
|  |  | MP only – Planting only | -0.377 | 0.373 | 34.6 | -1.013 | 0.7433 | EMMs |
|  |  | MP only – Replanted palms | -0.149 | 0.373 | 34.6 | -0.399 | 0.9781 | EMMs |
|  |  | Planting only – Replanted palms | 0.229 | 0.385 | 34.6 | 0.594 | 0.9333 | EMMs |
| Metric | Type | Comparison | Estimate | SE | df | t-ratio | p-value | Test |
| Leaf Litter (~5 cm) | Max | Planting + MP – MP only | -2.030 | 0.674 | 18 | -3.012 | 0.0344 | EMMs |
|  |  | Planting + MP – Planting only | -1.227 | 0.674 | 18 | -1.821 | 0.2962 | EMMs |
|  |  | Planting + MP – Replanted palms | -0.258 | 0.674 | 18 | -0.382 | 0.9804 | EMMs |
|  |  | MP only – Planting only | 0.803 | 0.674 | 18 | 1.191 | 0.6398 | EMMs |
|  |  | MP only – Replanted palms | 1.773 | 0.674 | 18 | 2.630 | 0.0734 | EMMs |
|  |  | Planting only – Replanted palms | 0.970 | 0.674 | 18 | 1.439 | 0.4928 | EMMs |
|  | Mean | Planting + MP – MP only | 0.448 | 0.460 | 19.6 | 0.974 | 0.7654 | EMMs |
|  |  | Planting + MP – Planting only | -0.602 | 0.426 | 19.6 | -1.412 | 0.5068 | EMMs |
|  |  | Planting + MP – Replanted palms | -0.930 | 0.460 | 19.6 | -2.020 | 0.2147 | EMMs |
|  |  | MP only – Planting only | -1.050 | 0.460 | 19.6 | -2.282 | 0.1366 | EMMs |
|  |  | MP only – Replanted palms | -1.378 | 0.492 | 19.6 | -2.801 | 0.0503 | EMMs |
|  |  | Planting only – Replanted palms | -0.328 | 0.460 | 19.6 | -0.712 | 0.8911 | EMMs |
|  | Minimum | Planting + MP – MP only | 1.058 | 0.417 | 19.6 | 2.538 | 0.0847 | EMMs |
|  |  | Planting + MP – Planting only | -0.333 | 0.386 | 19.6 | -0.864 | 0.8233 | EMMs |
|  |  | Planting + MP – Replanted palms | 0.414 | 0.417 | 19.6 | 0.993 | 0.7552 | EMMs |
|  |  | MP only – Planting only | -1.392 | 0.417 | 19.6 | -3.338 | 0.0162 | EMMs |
|  |  | MP only – Replanted palms | -0.644 | 0.446 | 19.6 | -1.446 | 0.4873 | EMMs |
|  |  | Planting only – Replanted palms | 0.747 | 0.417 | 19.6 | 1.792 | 0.3066 | EMMs |

**Supplementary Table S5:** Post Hoc comparisons for Vapour Pressure Deficit between restoration treatments between treatments (Planting + MP: enrichment planting with mature oil palm; MP only: mature oil palm only; Planting only: mature oil palm removed with enrichment planting; Palms replanted: control— oil palms planted to river edge) and for both field seasons (Dry; 2023 & Wet; 2024).

| Post Hoc VPD comparison | | | | | | | |
| --- | --- | --- | --- | --- | --- | --- | --- |
| Dry Season (2023: July) | | | | | | | |
| Metric | Comparison | Estimate | SE | df | t. ratio | p-value | Test |
| Max | Planting + MP – MP only | -5.003 | 1.94 | 24.1 | -2.579 | 0.0728 | EMMs |
|  | Planting + MP – Planting only | -4.198 | 1.86 | 24.1 | -2.260 | 0.1358 | EMMs |
|  | Planting + MP – Replanted palms | -7.771 | 2.24 | 24.1 | -3.469 | 0.0100 | EMMs |
|  | MP only – Planting only | 0.804 | 1.86 | 24.1 | 0.433 | 0.9722 | EMMs |
|  | MP only – Replanted palms | -2.769 | 2.24 | 24.1 | -1.236 | 0.6108 | EMMs |
|  | Planting only – Replanted palms | -3.573 | 2.17 | 24.1 | -1.647 | 0.3724 | EMMs |
| Mean | Planting + MP – MP only | -1.183 | 0.433 | 24.1 | -2.735 | 0.0525 | EMMs |
|  | Planting + MP – Planting only | -0.807 | 0.414 | 24.1 | -1.947 | 0.2358 | EMMs |
|  | Planting + MP – Replanted palms | -2.028 | 0.500 | 24.1 | -4.058 | 0.0024 | EMMs |
|  | MP only – Planting only | 0.377 | 0.414 | 24.1 | 0.909 | 0.8000 | EMMs |
|  | MP only – Replanted palms | -0.844 | 0.500 | 24.1 | -1.690 | 0.3506 | EMMs |
|  | Planting only – Replanted palms | -1.221 | 0.484 | 24.1 | -2.524 | 0.0813 | EMMs |
| Wet Season (2024: February) | | | | | | | |
| Metric | Comparison | Estimate | SE | df | t. ratio | p-value | Test |
| Max | Planting + MP – MP only | -0.648 | 0.325 | 26.9 | -1.996 | 0.2144 | EMMs |
|  | Planting + MP – Planting only | -0.974 | 0.309 | 26.9 | -3.157 | 0.0192 | EMMs |
|  | Planting + MP – Replanted palms | -1.001 | 0.348 | 26.9 | -2.880 | 0.0364 | EMMs |
|  | MP only – Planting only | -0.326 | 0.336 | 26.9 | -0.971 | 0.7671 | EMMs |
|  | MP only – Replanted palms | -0.353 | 0.372 | 26.9 | -0.949 | 0.7791 | EMMs |
|  | Planting only – Replanted palms | -0.027 | 0.358 | 26.9 | -0.075 | 0.9998 | EMMs |
| Mean | Planting + MP – MP only | -0.648 | 0.325 | 26.9 | -1.996 | 0.2144 | EMMs |
|  | Planting + MP – Planting only | -0.974 | 0.309 | 26.9 | -3.157 | 0.0192 | EMMs |
|  | Planting + MP – Replanted palms | -1.001 | 0.348 | 26.9 | -2.880 | 0.0364 | EMMs |
|  | MP only – Planting only | -0.326 | 0.336 | 26.9 | -0.971 | 0.7671 | EMMs |
|  | MP only – Replanted palms | 0.353 | 0.372 | 26.9 | -0.949 | 0.7791 | EMMs |
|  | Planting only – Replanted palms | -0.027 | 0.358 | 26.9 | 0.075 | 0.9998 | EMMs |

**Supplementary Table S6:** Post Hoc comparisons for the mean Diurnal Temperature range between restoration treatments between treatments (Planting + MP: enrichment planting with mature oil palm; MP only: mature oil palm only; Planting only: mature oil palm removed with enrichment planting; Palms replanted: control— oil palms planted to river edge).

| Post Hoc Comparisons for Diurnal Temperature Range | | | | | | | |
| --- | --- | --- | --- | --- | --- | --- | --- |
| Metric | Comparison | Estimate | SE | df | t. ratio | p-value | Test |
| Macroclimate (~1.5 m aboveground) | Planting + MP – MP only | -2.25 | 1.25 | 35.6 | -1.797 | 0.2915 | EMMs |
|  | Planting + MP – Planting only | -1.04 | 1.25 | 35.6 | -0.829 | 0.8403 | EMMs |
|  | Planting + MP – Replanted palms | -4.60 | 1.30 | 35.6 | -3.543 | 0.0059 | EMMs |
|  | MP only – Planting only | 1.21 | 1.25 | 35.6 | 0.968 | 0.7684 | EMMs |
|  | MP only – Replanted palms | -2.34 | 1.30 | 35.6 | -1.807 | 0.2868 | EMMs |
|  | Planting only – Replanted palms | -3.56 | 1.30 | 35.6 | -2.742 | 0.0448 | EMMs |
| Soil | Planting + MP – MP only | 0.787 | 0.937 | 34.6 | 0.841 | 0.8347 | EMMs |
|  | Planting + MP – Planting only | 1.176 | 0.970 | 34.6 | 1.213 | 0.6230 | EMMs |
|  | Planting + MP – Replanted palms | 0.119 | 0.970 | 34.6 | 0.123 | 0.9993 | EMMs |
|  | MP only – Planting only | 0.389 | 0.970 | 34.6 | 0.401 | 0.9779 | EMMs |
|  | MP only – Replanted palms | -0.668 | 0.970 | 34.6 | -0.689 | 0.9005 | EMMs |
|  | Planting only – Replanted palms | -1.057 | 1 | 34.6 | -1.056 | 0.7183 | EMMs |
| Leaf Litter | Planting + MP – MP only | -0.900 | 1.14 | 19.6 | -0.793 | 0.8569 | EMMs |
|  | Planting + MP – Planting only | -0.692 | 1.05 | 19.6 | -0.658 | 0.9114 | EMMs |
|  | Planting + MP – Replanted palms | -4.267 | 1.14 | 19.6 | -3.758 | 0.0064 | EMMs |
|  | MP only – Planting only | 0.208 | 1.14 | 19.6 | 0.184 | 0.9977 | EMMs |
|  | MP only – Replanted palms | -3.367 | 1.21 | 19.6 | -2.774 | 0.0531 | EMMs |
|  | Planting only – Replanted palms | -3.575 | 1.14 | 19.6 | -3.149 | 0.0243 | EMMs |

**Supplementary table S7:** Summary of linear model results assessing the influence of vegetation structure (PC1) and restoration treatment (Planting + MP: enrichment planting with mature oil palm; MP only: mature oil palm only; Planting only: mature oil palm removed with enrichment planting; Palms replanted: control— oil palms planted to river edge) on microclimate variables (Maximum, Mean, Minimum) from Linear Model (ANCOVA).

| Season | Temperature Variable | Statistical Test | PC1 Effect (p) | Treatment Effect (p) | PC1 × Treatment Interaction (p) | Residual Normality (p) | Homogeneity of Variance (p) |
| --- | --- | --- | --- | --- | --- | --- | --- |
| Wet | Maximum temperature | Linear model | 0.031 | 0.049 | 0.127 | 0.360 | 0.147 |
| Dry | Maximum temperature | Linear model | <0.001 | 0.664 | 0.214 | 0.717 | 0.151 |
| Dry | Mean temperature | Linear model | <0.001 | 0.901 | 0.222 | 0.640 | 0.173 |
| Wet | Mean temperature | Linear model | 0.156 | 0.085 | 0.334 | 0.919 | 0.358 |
| Wet | Minimum temperature | Linear model | 0.156 | 0.085 | 0.334 | 0.919 | 0.358 |
| Dry | Minimum temperature | Linear model | <0.001 | 0.140 | 0.950 | 0.672 | 0.172 |

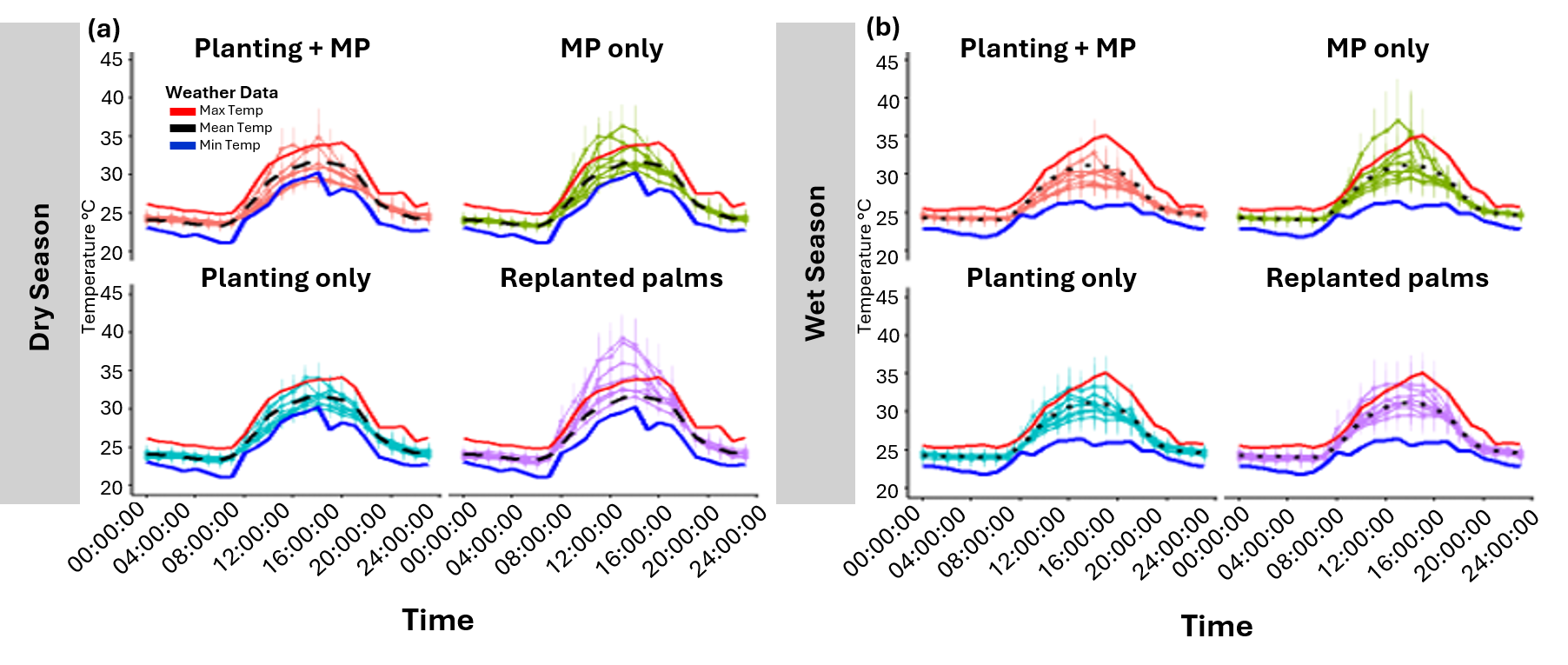

**Figure 6.** **Variations in microclimate temperature range.** Hourly microclimates (**a & b**) were measured across restoration treatments (Planting + MP: enrichment planting with mature oil palm; MP only: mature oil palm only; Planting only: mature oil palm removed with enrichment planting; Palms replanted: control— oil palms planted to river edge), within the margin and compared to nearby weather stations for both field seasons. Data from Dry **(a)** and Wet **(b)** were compared to nearby weather station data outside the riparian margin, which is represented by overlaid lines showing maximum (red), mean (black), and minimum (blue) temperatures. Hourly temperatures were averaged each hour.

**Supplementary table S8:** PCA intraset correlation values (Very Strong Correlations ≥ 0.90 in bold).

|  | Dim.1 | Dim.2 | Dim.3 | Dim.4 | Dim.5 |
| --- | --- | --- | --- | --- | --- |
| Canopy.Height | **0.923037** | -0.23778 | 0.203641 | -0.22358 | 0.002047 |
| MeanFrac | -0.18991 | **0.96866** | 0.114459 | -0.04549 | 0.102289 |
| ENL | 0.853953 | -0.48325 | 0.025717 | 0.173513 | 0.080395 |
| SSCI | 0.602212 | 0.745303 | 0.247297 | 0.123349 | -0.07412 |
| Canopy.Openness | -0.72689 | -0.50527 | 0.463778 | 0.034005 | 0.008919 |
